# A decomposition of a phylogenetically-informed distance into basal and terminal components

**DOI:** 10.1101/2025.04.17.649379

**Authors:** Julia Fukuyama

## Abstract

Ecologists needing to quantify differences between communities of organisms often use measures of dissimilarity that incorporate both differences in species composition and information about the phylogenetic relatedness of the species. Many variants on these distances are available to analysts, but their properties are not well developed. We analyze a phylogenetically-informed distance that has been described many times in the literature under different names. We show that we can decompose this distance into pieces that describe basal and terminal phylogenetic structure and show that it places an overwhelming amount of weight on the basal phylogenetic structure. We show that a related class of distances can be interpreted as modulating the influence of the basal structure, how this modification can give more power for identifying different scales of phylogenetically-structured effects, and show examples in simulated and real datasets.

## 1 Introduction

Ecologists have long been interested in quantifying dissimilarities between communities of organisms [5]. Many of the classic measures for this task, such as the Jaccard Index [25] or the Bray-Curtis dissimilarity [7], operate under the implicit assumption that all the species are equally distinct. Since species are not in fact equally distinct from each other, more nuanced measures have been introduced that incorporate information about the similarities between the species in these communities [43, 11, 36]. These similarities can in principle be anything, but a common metric for species similarity in this context is phylogenetic relatedness [47, 35].

As is well understood in the literature, no one measure can describe every possible aspect of the data. For instance, measures of alpha diversity can vary in how much weight they place on rare versus abundant species [27]. Since there is not generally going to be a clear choice for the correct amount of weight to put on rare species, some in the literature advocate computing diversity profiles that give the diversity measure for the entire range of possible weights [28]. In the context of phylogenetically-informed alpha- and beta-diversity measures, there have been empirical demonstrations that these measures can tend to emphasize either terminal or basal phylogenetic diversity [32, 45, 26], along with recommendations to use both kinds of measure so as to get a better understanding of the data [26]. However, these have been empirical demonstrations, a mathematical explanation of this phenomenon is not, to our knowledge, available.

In this paper, we help fill this gap by providing a mathematical explanation of why one phylogenetically-informed measure of beta diversity or dissimilarity emphasizes basal phylogenetic structure. The measure we focus on is derived from Rao’s axiomatization of diversity, and therefore it or something very similar has been widely used for measuring beta diversity [12, 47, 24] and in ordination [37].

Our results make heavy use of the idea of “derived variables” that are constructed from the abundance or relative abundance profiles and the phylogenetic tree. Instead of thinking of the distance as being between vectors giving the abundance of each species at a particular site, we think of the distance as being between vectors containing the values for a new set of variables that are derived from a combination of the abundance profiles and the phylogenetic tree. We will see that these derived variables lie on a gradient ranging from variables related to the basal phylogenetic structure to variables related to the terminal phylogenetic structure. These variables are similar in spirit to the derived variables used in other phylogenetically-informed distances, where the derived variables are the abundances or relative abundances of species that descend from each node in the tree (see [35] and the references therein for examples). In that case, we would say that derived variables that describe the relative abundances of large clades are related to the basal structure of the tree and derived variables that describe the relative abundances of small clades are related to the terminal structure of the tree.

Our results show that a small number of these derived variables tend to have much more weight than the others. These highly-weighted derived variables will tend to be those that are related to the basal phylogenetic structure. In some cases, this is almost as if the abundance profiles had been aggregated to the phylum level and dissimilarities computed based on the aggregated data. We suspect that this is not always the desired behavior of a phylogenetically-informed dissimilarity measure, and we therefore show how one can use a family of distances already in the literature to get a profile of distances, similar to Leinster and Cobbold’s diversity profiles, that progressively equalize the weights given to these derived variables, in effect interpolating between the original phylogenetically-informed distance and a distance that treats all the species as being equally distinct. We also show that downweighting the basal phylogenetic features can give increased statistical power for detecting certain kinds of differences between communities.

The format of the remainder of the paper is as follows: As there are a variety of distances in the literature that are identical to or similar to the one we study here, we first review these distances. We also give some mathematical background that is necessary for our technical results. We show how to rewrite these distances as a weighted Euclidean distance on a set of derived variables. We give explicit expressions for these derived variables and the weights assigned to them for two classes of trees. These expressions are, to our knowledge, new. We show that a previously-described family of distances allows us to modulate the weight accorded to the basal phylogenetic structure and show that different members of this family are better at detecting effects at different phylogenetic scales. These power results are new. Finally, we show on real and simulated datasets how this family of distances can be used to gain more insight into community structure.

## 2 Background

The phylogenetically-informed distance we focus on does not have a consistent name in the literature. A special case of its square is called *H* in [12], its square is called Rao’s DISC in [47], it is called *δ*^*CO*^ in the context of a method called double principal coordinates analysis (DPCoA) [37], and a normalized version is called *P*_*ST*_ in [24]. This distance is commonly used for microbiome data analysis in the context of DPCoA [41, 16, 49, 6, 22] and is implemented in several of the standard packages for analyzing ecological [15] and microbiome [33] data. In addition to being exactly the same as several other distances that have been described before, this distance is closely related to nearly any measure that is based on Rao’s axiomatization of diversity [43]. It is also closely related [20, 21] to weighted UniFrac [30], another popular phylogenetically-informed distance. See Section A.1 for a description of the other measure that correspond exactly to the one studied here, and a review of these and other related measures in [47]. We will call this distance the PQ distance because it is a <u>p</u>hylogenetically-informed distance based on Rao’s <u>q</u>uadratic entropy.

Before defining this distance, we review Rao’s axiomatization of diversity and define a similarity matrix **T** which will be of central importance.

### 2.1 Rao’s diversity and dissimilarity coefficients

Rao defined diversity and dissimilarity coefficients [43] as follows. Suppose we have *p* species, and **Δ** is a *p × p* matrix whose *ij*th element, *δ*_*ij*_, gives the dissimilarity between species *i* and species *j*. **Δ** being a dissimilarity matrix means that *δ*_*ii*_ = 0 and *δ*_*ij*_ ≥ 0 for all *i* ≠ *j*. Then if **x** is a *p*-vector whose *i*th element, *x*_*i*_, gives the relative abundance of the *i*th species, Rao’s diversity coefficient is

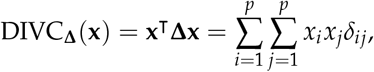

and his dissimilarity coefficient is

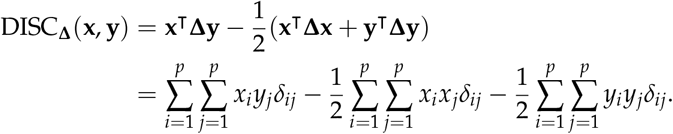

When **x** and **y** lie in the simplex, DIVC_**Δ**_ can be interpreted as the average dissimilarity between randomly selected pairs in a given sample, **x**^⊺^**Δy** can be interpreted as the average dissimilarity between a randomly selected member of **x** and a randomly selected member of **y**, and DISC_**Δ**_ can be interpreted as the excess expected dissimilarity associated with picking one member of the pair from **x** and the other from **y** vs. picking both members of the pair from **x** or both members of the pair from **y**.

From the definition, we can easily see that, so long as the diagonal of **Δ** is zero and all the other elements are non-negative, the smallest value that DIVC_**Δ**_(**x**) can take for **x** a relative abundance vector is 0 when **x** places all its mass on one species. The maximizing value for general **Δ** matrices is more complicated; for a discussion see [38]. Later, we will specialize to **Δ** encoding dissimilarities related to the phylogenetic tree, in which case we will be able to better describe the maximizing and minimizing values.

### 2.2 The similarity matrix T

An object of central importance to this manuscript is a *p × p* matrix we will call **T**. Given a rooted tree with *p* leaves, the corresponding matrix **T** has *ij*th element *t*_*ij*_ such that *t*_*ij*_ equal to the distance between the root and the most recent common ancestor of the *i*th and *j*th leaves. The diagonal elements *t*_*ii*_ are the distance between the root and the *i*th leaf. Therefore, the elements of **T** vary between 0 (for pairs of leaves on opposite sides of the root) and the height of the tree. If the tree is ultrametric, the distances from any of the leaves to the root are all equal and therefore all the diagonal elements of **T** will be equal to each other and equal to the height of the tree. For any *i* ≠ *j*, we always have *t*_*ij*_ *< t*_*ii*_, and the maximal value of **T** is the distance between the root and the leaf that is farthest from the root. Particularly if the tree is ultrametric, **T** can be interpreted as a similarity matrix, with *t*_*ij*_ describing how similar leaves *i* and *j* are according to the tree.

In addition to being a similarity matrix, this matrix is the covariance matrix of a trait that has undergone a Brownian motion along the tree [9]. It was used by Cavalli-Sforza and his collaborators in analyzing human populations [9], by Felsenstein (who also calls it **T**) in his method for maximum likelihood estimation of phylogenetic trees based on quantitative characteristics [17], and has been consistently used for testing or controlling for phylogenetic effects ever since. For instance, Felsenstein also uses the Brownian motion model and the covariance matrix implied by it in developing his method of phylogenetically independent contrasts [18]. **T** and the Brownian motion model that gives rise to it is also used in testing for phylogenetic signal in a trait [44] (referred to in that paper as **C**). The papers described above are highly influential but older — for a representative example of a recent paper that builds on the same model and covariance structure, see [4].

### 2.3 The PQ distance

With **T** defined, the definition of the PQ distance is as follows: Suppose **x** and **y** are *p*-vectors whose *i*th element measures the amount of the *i*th species in a community. We will start off assuming that these are relative abundance vectors (i.e., all their elements are non-negative and the sum of their elements is 1), but see the last paragraph of Section A.5 for why one might want to used some transformation of relative abundances. Furthermore, assume that **T** is a *p × p* matrix corresponding to a phylogenetic tree with *p* leaves as defined in the previous section. Then the PQ distance between **x** and **y** is

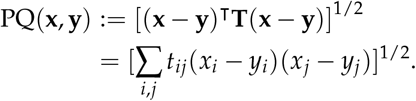

The matrix form of this equation will be useful later. In the results section, we will look in more detail at **T**’s properties and show how they can help us interpret this distance.

The scalar/sum version is of the definition is not very transparent, but we can gain some intuition by breaking the sum down into pieces. Focusing first on the elements of the sum for which *i* = *j*, we see that the larger the difference in relative abundance of a species between **x** and **y** (if (*x*_*i*_ − *y*_*i*_)^2^*t*_*ii*_ is large), the larger PQ(**x, y**). The elements of the sum for which *i* ≠ *j* make the PQ distance larger to the extent that species *i* and *j* are similar to each other (*t*_*ij*_ is large) and to the extent that they are either both over-represented in **x** compared to **y** or both under-represented in **x** compared to **y** ((*x*_*i*_ *− y*_*i*_)(*x*_*j*_ − *y*_*j*_) is positive). On the other hand, if species *i* is over-represented in **x** compared to **y** and species *j* is under-represented in **x** compared to **y** (that is, if (*x*_*i*_ *− y*_*i*_)(*x*_*j*_ *− y*_*j*_) is negative), that will make the PQ distance smaller, and it will make the distance *much* smaller if *i* and *j* are very similar (if *t*_*ij*_ is large).

If **x** and **y** are vectors containing relative abundances, the PQ distance can also be thought of as specializing DISC_**Δ**_ to the case where **Δ** gives the patristic distances (the length of the shortest path on the tree) between the species represented by the tree. In that case, if we let diag(**T**) = (*t*_11_, … , *t*_*pp*_)^⊺^ and **1** be the *p*-vector with all entries equal to one, we can write **Δ** = diag(**T**)**1**^⊺^ + **1**diag(**T**)^⊺^ − 2**T**. The reason for this equality is that *δ*_*ij*_ is equal to the distance between *i* and the most recent common ancestor (MRCA) of *i* and *j* plus the distance between *j* and the MRCA of *i* and *j*. The distance between *i* and the MRCA of *i* and *j* is equal to *t*_*ii −*_ *t*_*ij*_, and the distance between *j* and the MRCA of *i* and *j* is equal to *t*_*jj*_ *− t*_*ij*_. Summing these gives that the patristic distance between *i* and *j* is *t*_*ii*_ + *t*_*jj*_ *−* 2*t*_*ij*_, and rewriting that expression in matrix form gives the expression above for **Δ**. Given that expression, we have

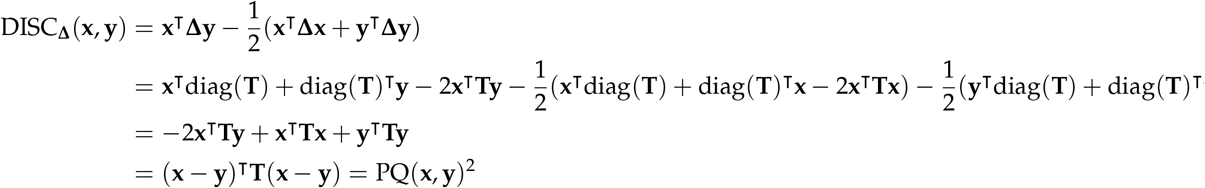

And so PQ(**x, y**) = [DISC_**Δ**_(**x, y**)]^1/2^ when **Δ** is the matrix containing patristic distances between the species. This has been shown before [41].

Because **T** is positive definite, the minimum value PQ can take is zero (for **x** = **y**). If **x** and **y** are relative abundance vectors, PQ(**x, y**) is maximized when **x** is the population consisting solely of leaf *i* and **y** is the population consisting solely of leaf *j* for *i* and *j* the leaves that are farthest apart on the tree (see Theorem 1 for proof). For ultrametric trees, *i* and *j* can be taken to be any pair of leaves on opposite sides of the root.

### 2.4 Derived variables

In this paper, we will differentiate between the measured variables, i.e., the measurements that were made when conducting the study, and derived variables, which are new variables that are constructed using the information in the measured variables. For instance, in many ecological studies, the measured variables are species counts, and we have one measured variable for each species. Derived variables can be something as simple as log-transforming a variable to reduce its skewness, but the derived variables we describe in this paper will combine information from multiple measured variables.

The sorts of derived variables we will be using here will be linear in either abundance or relative abundance. Suppose for each site we have a vector **x** = (*x*_1_, … , *x*_*p*_) giving either the abundance or the relative abundance of each of *p* species. Then if we have a *p*-dimensional vector **a** = (*a*_1_, … , *a*_*p*_), we can define a derived variable 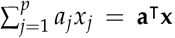. We will refer to these sorts of vectors as *coefficient vectors*, and each element of **a** is the coefficient for one species.

For most coefficient vectors **a**, the derived variable 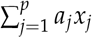 will not be meaningful. However, in certain cases they are interpretable. Consider the following examples:

1. Suppose **x** contains the abundances of the *p* species at a given site. Then if *a*_1_ = · · · = *a*_*p*_ = 1, then the feature 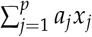 gives the overall abundance at that site.
2. Suppose **x** contains the abundances of the *p* species at a given site, and we are interested in a particular phylum that has *m* species. If *a*_*j*_ = 1/*m* for all *j* such that the *j*th species is a member of the phylum of interest, the feature 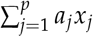 would be the average abundance of species in that phylum.
3. Suppose **x** contains relative abundances. If *a*_1_ = 1, *a*_2_ = −1, and the remaining *a*_*j*_’s are 0, the feature 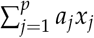 would be the difference between the relative abundance of species 1 and species 2.

Finally, notice that for any coefficient vector **a** and any positive number *s*, the coefficient vector *s***a** = (*sa*_1_, … , *sa*_*p*_) gives rise to a derived variable that has the same information as **a**, just at a different scale. Therefore, we will generally want to use normalized coefficient vectors, those that satisfy 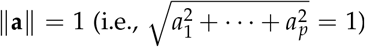. Given an unnormalized coefficient vector, we can always create a normalized coefficient vector by dividing the unnormalized coefficient vector by its norm.

These sorts of derived variables are central to our results, and we will be particularly interested in the relationship between the coefficient vectors and the phylogenetic tree. Some examples of these coefficient vectors (the unnormalized versions) are shown in Figure 1. Panels A-C of that figure show the coefficient vectors corresponding to the linear derived variables used by the PQ distance for three different trees. We will discuss these in more detail in the following sections, but for now, we will describe how to read the plot and give two examples.

**Figure 1:**
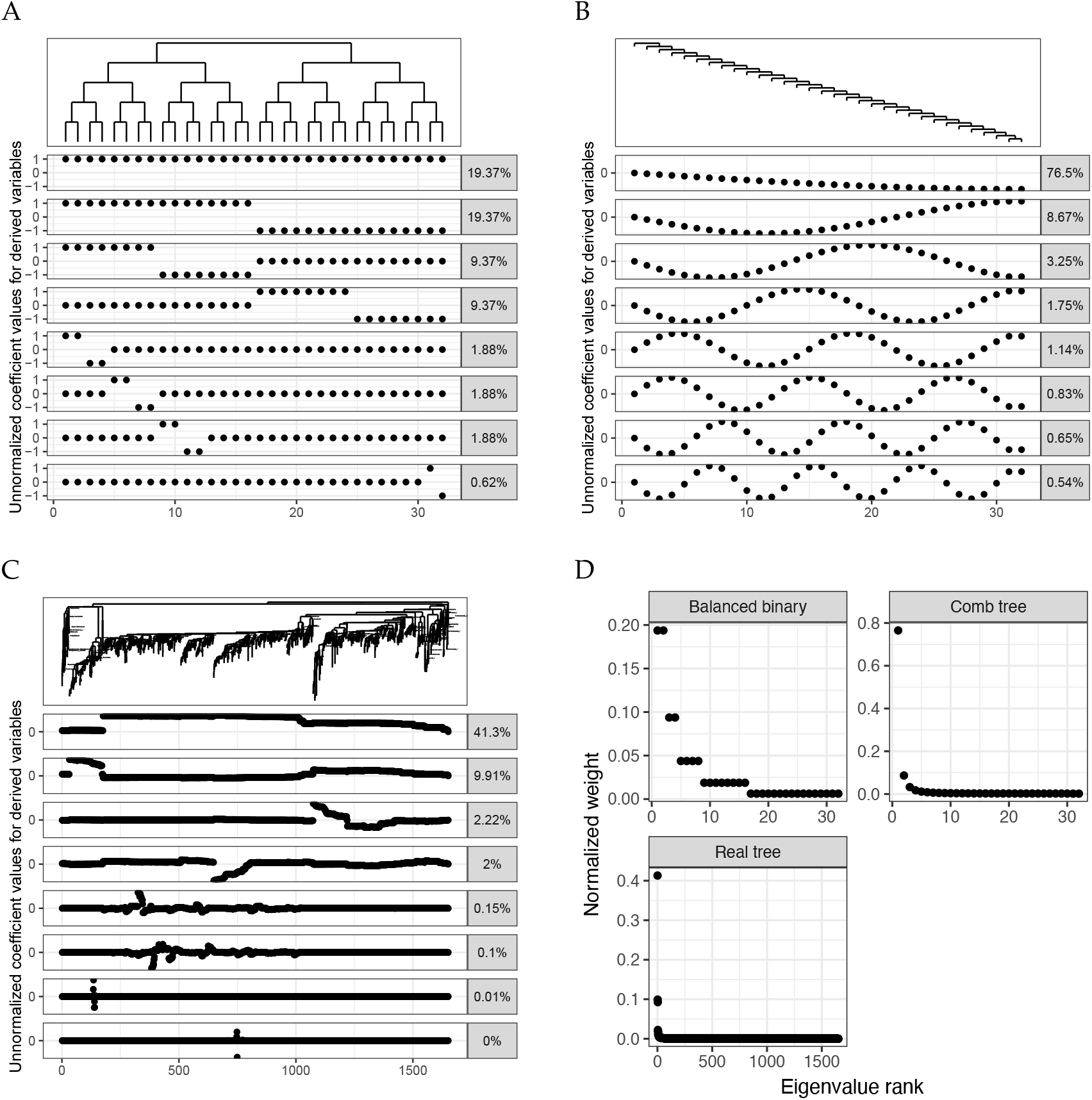
Coefficient values (A-C) and weights (D) for selected derived variables corresponding to the PQ distance with different tree topologies. (A-C): Coefficient values for derived variables used for the PQ distance with (A) the perfect binary tree with 32 leaves (B) the comb tree with 32 leaves and (C) a real tree taken from a microbiome data set. In each panel, the tree is plotted in the top section and coefficients for selected derived variables are plotted in the bottom section. Below the tree, each facet corresponds to one derived variable. Within the facet, each point corresponds to the coefficient for a species, and the points are aligned to the leaves of the corresponding tree. Each facet is labeled with the fraction of the total weight that is assigned to the derived variable represented in the facet. The coefficient vectors would be normalized prior to being used in the PQ distance, so only the signs and relative magnitudes within each row matter. (D): The distribution of weights for all the derived variables for the trees shown in panels (A-C). Each point corresponds to the weight for one derived variable, and the weights are plotted in decreasing order. The weights are normalized so that the sum of the weights for each tree is equal to 1.

Figure 1A gives examples of the coefficients for linear derived variables for the perfect binary tree with 32 leaves. The top panel of Figure 1A shows the perfect binary tree with 32 leaves. Each row below the tree plots one example of a coefficient vector **a**. The coefficients for each of the 32 species, *a*_1_, *a*_2_, … , *a*_32_ are plotted from left to right within a row, and they are aligned under the tree so that the value *a*_1_ is directly under leaf 1, the value *a*_2_ is directly under leaf 2, and so on. Notice that the first and last rows of Figure 1A (labeled “19.37%” and “.62%”) correspond to the sort of derived variables described in (1) and (3) above. In the top row “19.37%” we see that all the *a*_*j*_ values are equal to 1, and so the corresponding derived variable corresponds to an overall abundance. The bottom row (labeled “.62%”) has the *a*_*j*_ values equal to 0 for all but two variables. Those two variables are sister species, one of which has weight 1 and the other of which has weight −1. The derived variable corresponding to the coefficients in that row therefore describes the difference in abundance or relative abundance between the two sister species.

### 2.5 Weighted Euclidean distance

Suppose we have two *p*-dimensional vectors **x** and **y**, and a set of *p* weights *w*_1_, … , *w*_*p*_. The weighted Euclidean distance between **x** and **y** with weights *w*_1_, … , *w*_*p*_ is 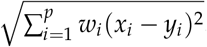. The interpretation is that some of the variables are more important than others, and the degree of importance of variable *i* is encoded in the weight *w*_*i*_.

### 2.6 Eigendecomposition

One of the mathematical tools we will use here is the eigendecomposition. For any symmetric positive definite matrix **M**, we can write **M** = **VDV**^⊺^, where **D** is a *p* × *p* diagonal matrix, and **V** is a *p* × *p* orthogonal matrix. We usually think of **V** as consisting of *p* column vectors concatenated together, **V** = (**v**^(1)^ · · · **v**^(*p*))^ , and each of the **v**^(*i*)^’s is called an *eigenvector*. **V** being an orthogonal matrix means that each of the **v**^(*i*)^’s has norm one, and each pair **v**^(*i*)^, **v**^(*j*)^ is orthogonal. The first property means that each **v**^(*i*)^ can be used as a normalized coefficient vector as described in Section 2.4. This also means that if we have a vector of measured variables **x**, 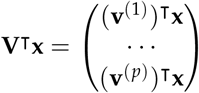 will be a vector of derived variables. The first derived variable will be a derived variable with coefficients given by **v**^(1)^, the second will be a derived variable with coefficients given by **v**^(2)^, and so on.

The values on the diagonal of **D**, *d*_11_, … , *d*_*pp*_ are called the eigenvalues of **M**.

The eigendecomposition is used in many contexts. One of the most familiar is PCA, where we eigendecompose the empirical covariance matrix to get the principal components. In this paper, we will use it in a different way. The eigendecomposition of **T** will allow us to decompose the PQ distance into basal and terminal pieces. Each eigenvector will give the coefficients for the type of linear derived variables described above, and each eigenvalue will be a weight in a weighted Euclidean distance as described in Section 2.5.

### 2.7 MPQ_*r*_ distances

The last definition we review is a class of distances that can be written in a similar way the same way as the PQ distance (used in [50]). The class of distances has a tuning parameter, *r*, and a given member of the family will be called the <u>m</u>odulated <u>p</u>hylogenetically-informed distance based on Rao’s <u>q</u>uadratic entropy with parameter *r* (MPQ_*r*_). It is defined as

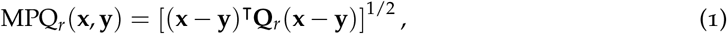

where

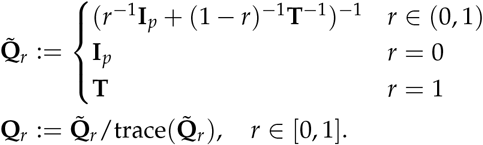

For an explanation of the scaling of **Q**_*r*_, see Section A.7.

## 3 Results

Our theoretical results are based on using the eigendecomposition of **T** to rewrite the PQ distance. Since eigendecomposition can be challenging to understand, we begin in Sections 3.1 and 3.2 by showing that for the perfect binary tree and the comb tree, the PQ distance can be rewritten as a weighted Euclidean distance on a set of derived variables and give a description of the derived variables and their weights. These results, which are new, follow from our derivation of the eigenvectors and eigenvalues of **T** for these specific trees. While the derived variables and weights introduced in these sections come directly from the eigendecomposition of **T**, we put off explicitly formulating the PQ distance in terms of eigendecomposition until Section 3.3.

### 3.1 Rewriting the PQ distance for the perfect binary tree

Suppose we have two vectors **x** and **y**, each of length *p*, that give the relative abundances of of each of *p* species at two sites. Suppose further that the phylogenetic relationships among the species are given by a perfect binary tree (i.e., a tree for which each non-leaf node has two children and all of the levels of the tree are completely filled) whose branch lengths are all 1. This means that *p* = 2^*K*^ for some positive integer *K*. We can define a distance between **x** and **y** in the following way:

1. Create vectors of derived variables 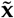 and 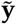. The first derived variable has unnormalized coefficient vector equal to (1, … , 1). The subsequent *p −* 1 derived variables correspond to each of the *p −* 1 nodes in the tree, and they have unnormalized coefficient vectors that take a value of 1 for all species descending from the left daughter of the node, a value of −1 for all species descending from the right daughter of the node, and 0 otherwise. In the notation of Section 2.4, the vector **a** giving the coefficients for the derived variable corresponding to a given node is

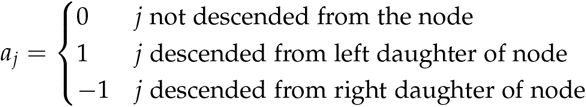
2. Give each node variable a weight based on how many descendants it has. A node variable with *d* descendants gets weight *d −* 1. This means that node variables corresponding to nodes with more descendants will have more weight. Give the first derived variable (corresponding to coefficient vector (1, … , 1)) a weight of *p −* 1 (the same weight as the most-highly-weighted node variable).
3. Compute a weighted Euclidean distance between 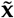 and 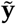 with the weights described in (2).

Note that since every time we go one level up the tree, the number of descendants of a node on that level doubles, the weights for the node variables approximately double every time we go one level up the tree. This means that for reasonably-sized trees, the top weights become much higher than the intermediate or small weights. Although this is a fairly steep weight progression, it is actually the most gradual we will see: both the comb tree and a real tree will have top weights that are even more extreme than the top weights for the perfect binary tree.

Also note that the derived variables described here have a simple interpretation. A derived variable corresponding to a node in the tree measures the difference in relative abundance between the descendants of the left and right daughters of that node. Therefore, a natural way to describe these derived variables is that some of them are more basal (those corresponding to nodes close to the root) and some are more terminal (those corresponding to nodes far from the root).

An example of these computations for the case of four species is given in Figure 2. In addition, examples of unnormalized coefficient vectors are given in Figure 1A. The top of Figure 1A shows a perfect binary tree with 32 leaves. Each row below the tree shows one unnormalized coefficient vector, and the row is labeled with the weight of the derived variable corresponding to that coefficient vector (as a percent of the total weight). Each point within a row corresponds to the coefficient for one species, and the points are positioned so that they are directly below the leaf in the tree they correspond to. So, for instance, the first row corresponds to the derived variable that gives equal weight to all the species, and has weight 19.37% of the overall weight. The second row corresponds to the derived variable that has coefficient values equal to 1 on the left side of the root and equal to − 1 on the right side of the root, and also has weight 19.37% of the overall weight. Subsequent rows correspond to the node variables described above for nodes that are increasingly far from the root, and therefore with weights that become smaller and smaller.

**Figure 2:**
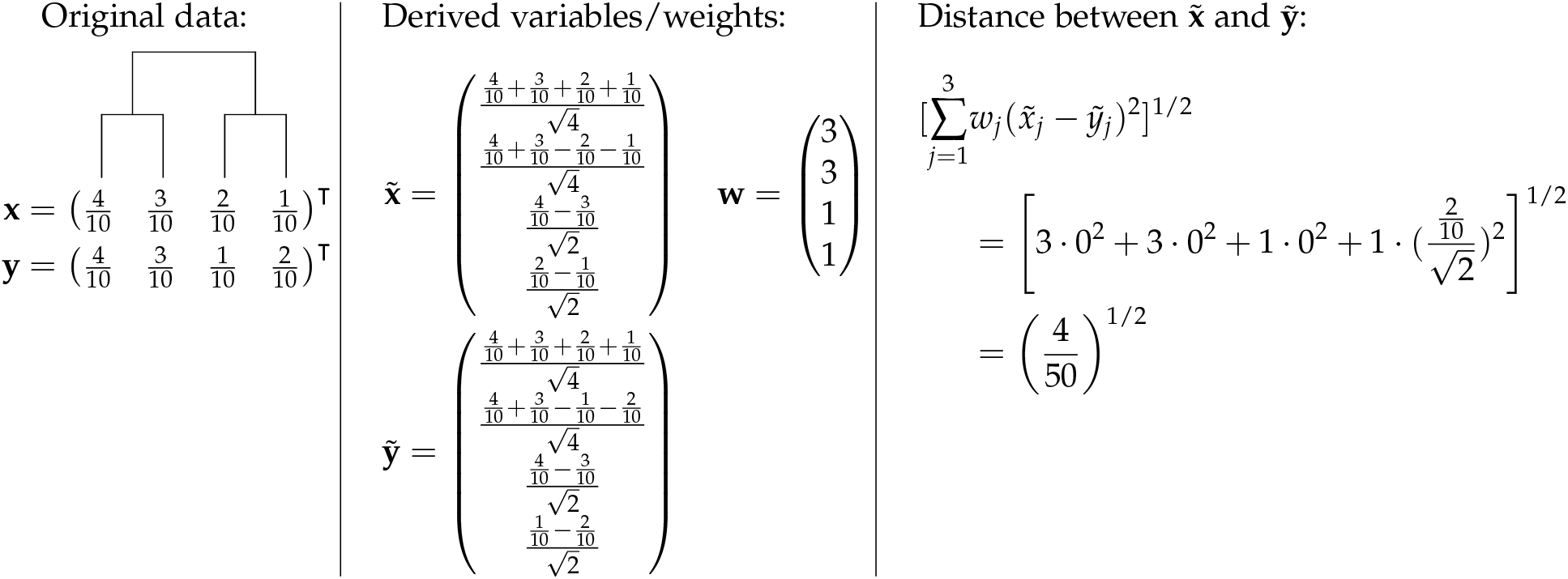
Example of computations to compute the distance between **x** and **y** defined in the text. Left panel is the tree and the relative abundances of the four species in **x** and **y**. Middle panel is 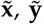, and **w**. For 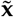 and 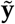, the first variable corresponds to an overall sum, the second variable corresponds to the root node, the second variable corresponds to the left daughter of the root, and the third variable corresponds to the right daughter of the root. Right panel shows the weighted Euclidean distance between 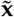 and 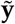with weights **w**.

It turns out that the weights and the coefficient vectors for the derived variables described here are the eigenvalues and eigenvectors of the matrix **T** for the perfect binary tree (see Theorem 2), and this procedure therefore gives the PQ distance (or the square root of Rao’s DISC_**Δ**_ or Chave and Chust’s *H* or *δ*^*CO*^). This reformulation gives another interpretation of these measures, at least when the species are related by a perfect binary tree.

This result raises two questions: first of all, can we write something similar for other kinds of trees? And second, if we think that the weights on the basal phylogenetic features are too high, is there anything we can do to mitigate that problem? As we will see in Sections 3.2 - 3.3, the answer to the first question is yes, although with less precision as the trees get more general. We will see one option for dealing with the second question in Section 3.5, whereby progressively diminishing the weight on the basal phylogenetic features will give us a family of distances that have differential weight on and sensitivity to the basal vs. terminal phylogenetic features.

### 3.2 Rewriting the PQ distance for the comb tree

The PQ distance does not have such a nice form for arbitrary trees, but we can approximately rewrite it in a similar way for the comb tree. The derived variables used for the construction of the distance will not have coefficients that correspond exactly to clades of different sizes as for the perfect binary tree, but they will still be features that are linear in the relative abundances.

The comb tree will be a tree for which the branches have equal length, and each internal node save the last (which has two leaf nodes as daughters) has one daughter which is a leaf and one daughter which is another internal node. An example is shown in the top panel of Figure 1B.

For this tree, the PQ distance is approximated by the following procedure, with the approximation approaching the exact distance as the number of species grows (See Theorem 3 for a precise statement and proof):

1. Number the species in ascending order of their distance to the root, so that species 1 is closest and species *p* is farthest.
2. Create *p* derived variables from the relative abundance vectors **x** and **y**. The unnormalized coefficient vector for the *j*th derived variable is 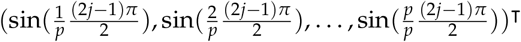.
3. Give the *j*th linear feature a weight of 4/[*π*^2^(2*j* − 1)^2^].
4. Compute the weighted Euclidean distance between the derived variables in (2) using the weights in (3).

As before, some of these coefficient vectors are plotted in Figure 1B. The comb tree is plotted at the top of that panel, and selected coefficient vectors are given below. The coefficient vectors are again labeled with the weight corresponding to that vector and they are plotted in decreasing order of weight. Comparing the coefficient vectors for the comb tree to the coefficient vectors for the perfect binary tree, we see differences in the particulars but some qualitative similarities. In both cases, the most highly-weighted derived variables have coefficient vectors have the largest-scale phylogenetic autocorrelation, i.e., the coefficient vectors have similar non-zero values over large portions of the tree. As we get to the less heavily-weighted variables, the coefficient vectors continue to show phylogenetic autocorrelation, but it is at smaller and smaller scales.

We also notice that the weight distribution is even more skewed for this set of variables than it was for the perfect binary tree. The top derived variable for the comb tree has weight equal to approximately 80% of the total, and this will be true regardless of the number of leaves in the tree.

The approximation results tell us that the quality of the approximation for the top eigenvectors (those for which the sine wave goes through ≪ *p* periods in the eigenvector) is quite good, but it can be very poor for the bottom eigenvectors (those for which the number of periods the sine wave goes through is a large fraction of *p*). This is essentially because for those eigenvectors the discretization error becomes very large. However, since the bottom eigenvectors have very small eigenvalues, they don’t harm the overall approximation very much. If we compute the correlation between the true matrix **T**^ct^ and the approximation **VDV**^*T*^ , where **V** is a *p × p* matrix whose columns are the approximate eigenvectors given in Theorem 3 and **D** is a *p × p* diagonal matrix whose elements are the approximate eigenvalues given in Theorem 3, the correlation between elements of the two vectors is .9988 when *p* = 20, .99998 when *p* = 100, and increases further with increasing *p*.

### 3.3 Rewriting the PQ distance for arbitrary trees

To understand how the results above were derived and to get analogous results for arbitrary trees, we need to rely on the eigendecomposition of **T**. This will allow us to rewrite the PQ distance as a weighted Euclidean distance between certain derived variables based on the species proportions or abundances and the phylogeny. The definition of these derived variables comes from the eigendecomposition of **T**, and so we will refer to them as the eigenvector features. If we write **T** = **VDV**^⊺^, where **V** is a *p × p* matrix whose columns are the eigenvectors of **T** and **D** is a *p × p* diagonal matrix whose diagonal elements are the eigenvalues corresponding to the eigenvectors in the columns of **V**, then we can write

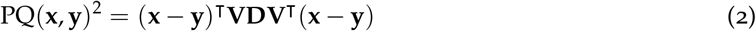

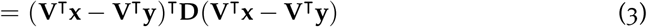

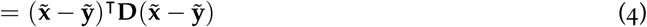

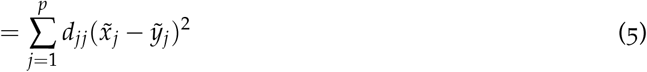

where 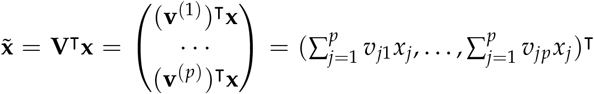 and *d*_*jj*_ is the *j*th diagonal element of **D**. We can interpret equation 5 as follows:

– 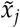is the value of the *j*th eigenvector feature (that is, the derived variable with coefficient vector **v**^(*j*)^) corresponding to the relative abundances in **x**, and similarly for 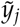.
– The PQ distance overall is a weighted Euclidean distance between the eigenvector features in 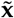 and 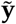. The weight associated with the *j*th eigenvector feature is *d*_*jj*_.

For this formulation of the PQ distance to be useful, we still need to understand the eigenvectors (columns of **V**) and eigenvalues (diagonal elements of **D**) of **T**. These quantities depend on the tree and are not generally available in closed form. The literature has an expression for trees with four leaves [9], and some limited results about the forms the eigenvectors can take for general trees [40]. Purdom’s thesis [40] contains the most extensive review of the known results about the eigenvalues and eigenvectors of **T**. She notes that all the authors who have studied this problem have noted that the eigenvectors of **T** have a sort of multiscale property, but that exact results are hard to come by.

The results in Sections 3.1 and 3.2 were derived from the fact that for the perfect binary tree, the eigenvectors of **T** were the coefficient vectors described in Section 3.1, and for the comb tree, the eigenvectors of **T** were approximately the sin functions defined in Section 3.2. For general trees we can compute these eigenvectors but not write them down in closed form. They tend to be qualitatively similar to the ones derived above for the comb tree and perfect binary tree, as we will see in the next section.

### 3.4 Derived variables and weights for a real tree

Next, we show that the eigenvector features for real trees are qualitatively similar to the eigen-vector features for the perfect binary tree and comb tree that we described in Sections 3.1 and 3.2. As this is a qualitative statement and not something mathematically precise, our evidence for this statement will be in the form of a visualization of the coefficients for the derived variables.

Recall that the coefficient vectors for a perfect binary tree and a comb tree are shown in Figure 1A/B. The main thing the coefficient vectors have in common is that the coefficient vectors with the highest weight are very smooth on the tree. That is, given the value of the coefficient vector for one species, the value of the coefficient vector for neighboring species is likely to be very similar. As the weights get smaller, this holds true for smaller and smaller neighborhoods. For the coefficient vectors with the very smallest weights, the opposite is true: if one species has a non-zero value for the coefficient vector, its closest neighbor is likely to have the opposite value for the coefficient vector.

The coefficient vectors for a real tree are similar in that regard. We see the coefficient vectors from a tree taken from a microbiome study [14], is shown in Figure 1C. As before, the coefficient vectors with the largest weight are smooth over large regions of the tree. As the weights get smaller, the coefficient vectors are smooth over smaller regions of the tree, and for the coefficient vectors with the smallest weights, the coefficients for neighboring species with non-zero weights tend to be the opposite of each other.

As with the comb tree, the eigenvalue spectrum/distribution of weights for the derived variables is very skewed, with the eigenvalue corresponding to the top eigenvector contributing over 40% of the total weight (Figure 1D). Since the tree in this dataset has about 1600 leaves and the same number of eigenvector features, the top eigenvector contributing 40% of the total weight not only means that the top feature contributes a large absolute amount of the weight, it also means that the ratio between the weight given to the top feature and subsequent features is extremely high. In the next section, we will see one way of modulating the high weight placed on the broad-scale/basal phylogenetic features.

### 3.5 MPQ_*r*_ distances use the same derived variables but with a less skewed weight distribution

Recall the MPQ_*r*_ distances, as defined in Equation (1), look very similar to the PQ distances, with the difference that **T** has been replaced with **Q**_*r*_. The definition of **Q**_*r*_ ensures that **Q**_*r*_ and **T** have the same eigenvectors, and they are in the same order. Therefore, the reasoning in Section 3.3 applies to the MPQ_*r*_ distances as well. They can also be rewritten as a weighted Euclidean distance on the same derived variables as the PQ distance can. The difference is in the weights put on those derived variables, which will tend to have a less skewed distribution than those used for the PQ distance.

When *r* = 1, the weights used in the MPQ_*r*_ distance will have the same relative magnitudes as the weights used in the PQ distance. When *r* = 0, the weights will all be equal, and **Q**_*r*_ is a scalar multiple of the identity matrix. This means that the MPQ_*r*_ distance with *r* = 0 is the standard Euclidean distance.

In between, as *r* decreases, the weights equalize gradually, and so the weight given to the top derived variable decreases relative to the weights given to the derived variables until we reach complete equality at *r* = 0. See Figure 3 for an illustration of the weights for a range of values of *r* for the case of a tree taken from a microbiome dataset.

**Figure 3:**
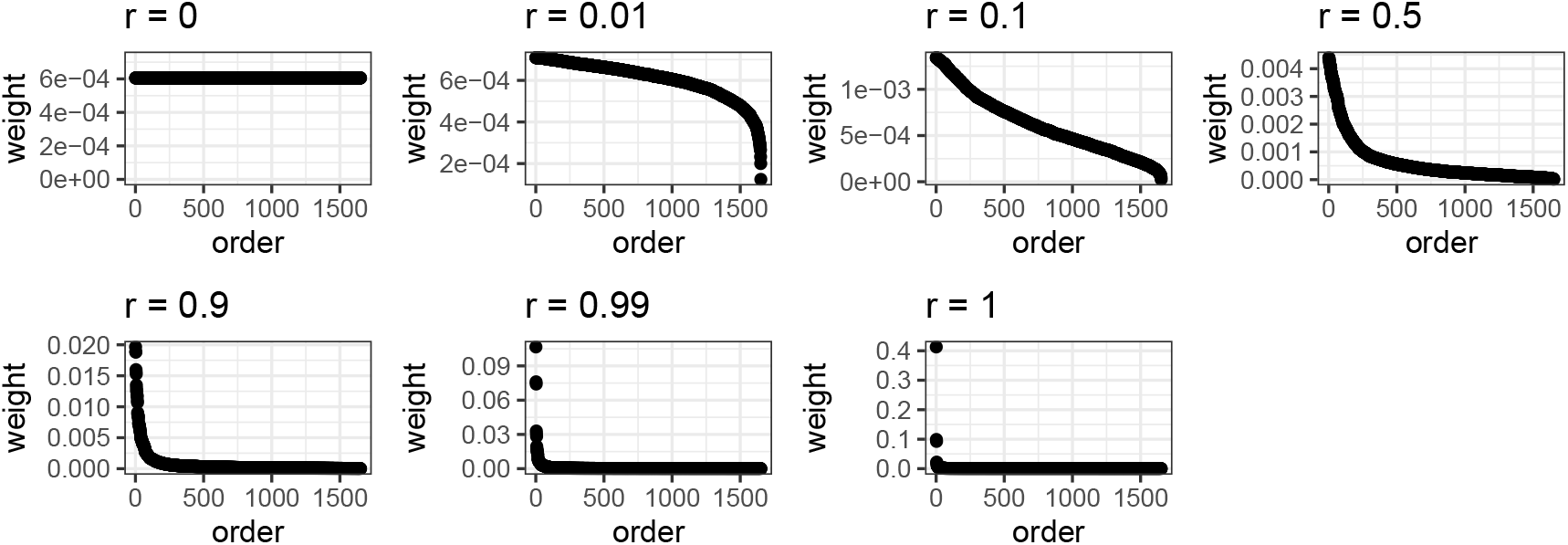
Weights of linear derived variables corresponding to a real tree for a range of values of *r* in MPQ_*r*_ distances. When *r* = 0, the weights are all equal. When *r* = 1, the weights are highly skewed, with the most-heavily-weighted linear derived variable being assigned more than 40% of the total weight. For intermediate values of *r*, the distribution of weights is more equal as *r* approaches 0 and is more skewed as *r* approaches 1.

To see that if the eigenvalue for the *j*th eigenvector of **T** is larger than the eigenvalue for the *i*th eigenvector of **T**, the same must be true for the *i*th and *j*th eigenvalues of **Q**_*r*_ for *r >* 0, let *d*_*i*_ and *d*_*j*_ denote the *i*th and *j*th eigenvalues of **T**. Then the *j*th and *i*th eigenvalues of 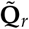 will be (*r*^−1^ + (1 − *r*)^−1^*d*_*j*_)^−1^ and (*r*^−1^ + (1 − *r*)^−1^*d*_*i*_)^−1^, and some simple rearrangement shows that (*r*^−1^ + (1 − *r*)^−1^*d*_*j*_)^−1^ *>* (*r*^−1^ + (1 − *r*)^−1^*d*_*i*_)^−1^ if *d*_*j*_ *> d*_*i*_.

### 3.6 MPQ_*r*_ distances allow for detection of different kinds of phylogenetically-structured effects

Earlier, we said that it seemed subjectively like the standard PQ distance puts too much weight on too small a number of features. There is a more rigorous sense in which the PQ distance does not use the ideal set of weights. The MPQ_*r*_ distance that allows us to see differences between communities the most clearly, both in the sense of what a plot of the distances looks like and in terms of statistical power, will tend to not be the PQ distance except in very specialized circumstances. The value of *r* that works the best will vary with the nature of the “true effect” – that is, the difference between two groups of communities, or a group of species that vary along a gradient – and with the nature of the noise. All else equal, larger values of *r* will tend to be better for effects that are related to the basal phylogenetic structure (e.g. differences in the abundance of large or old clades), and smaller values of *r* will tend to be better for effects that are related to the terminal phylogenetic structure (e.g. differences in the abundance of small or new clades).

The mathematical model that we can use to make this statement precise is the following: Suppose that **x, y** ∈ ℝ^*p*^ have distribution 𝒩 (***µ***_*x*_, Σ) and 𝒩 (***µ***_*y*_, Σ), respectively. Because of the measurement noise, even if the two communities have the same mean abundance, (that is, ***µ***_*x*_ = ***µ***_*y*_), MPQ_*r*_(**x, y**) will not be exactly zero. Rather, it will have a generalized chi squared distribution (see Section A.6 for a definition and the exact form of the distribution of MPQ_*r*_(**x, y**)). Therefore, in the standard statistical way, if we want to decide whether the observed value of the MPQ_*r*_ distance is large, we compare it to what we would expect to see if there was truly no difference between the two communities. If we fix ***µ***_*x*_, ***µ***_*y*_, and Σ, we can ask which value of *r* will tend to give the largest value of the MPQ_*r*_ distance relative to the noise (i.e., we can ask which value of *r* will have the most power in a test based on the MPQ_*r*_ distance).

The formal statistical test and exact expressions for its power as a function of ***µ***_*x*_, ***µ***_*y*_, and Σ are given in the Appendix (Section A.8). Using the expressions derived there, we can find numerically the values of *r* that maximize the power of the test. One example is shown in Figure 4A. In that case, we have 256 species connected to each other by a perfect binary tree whose branch lengths are all equal. We set Σ = **I**, and for each clade, we set

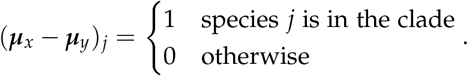

**Figure 4:**
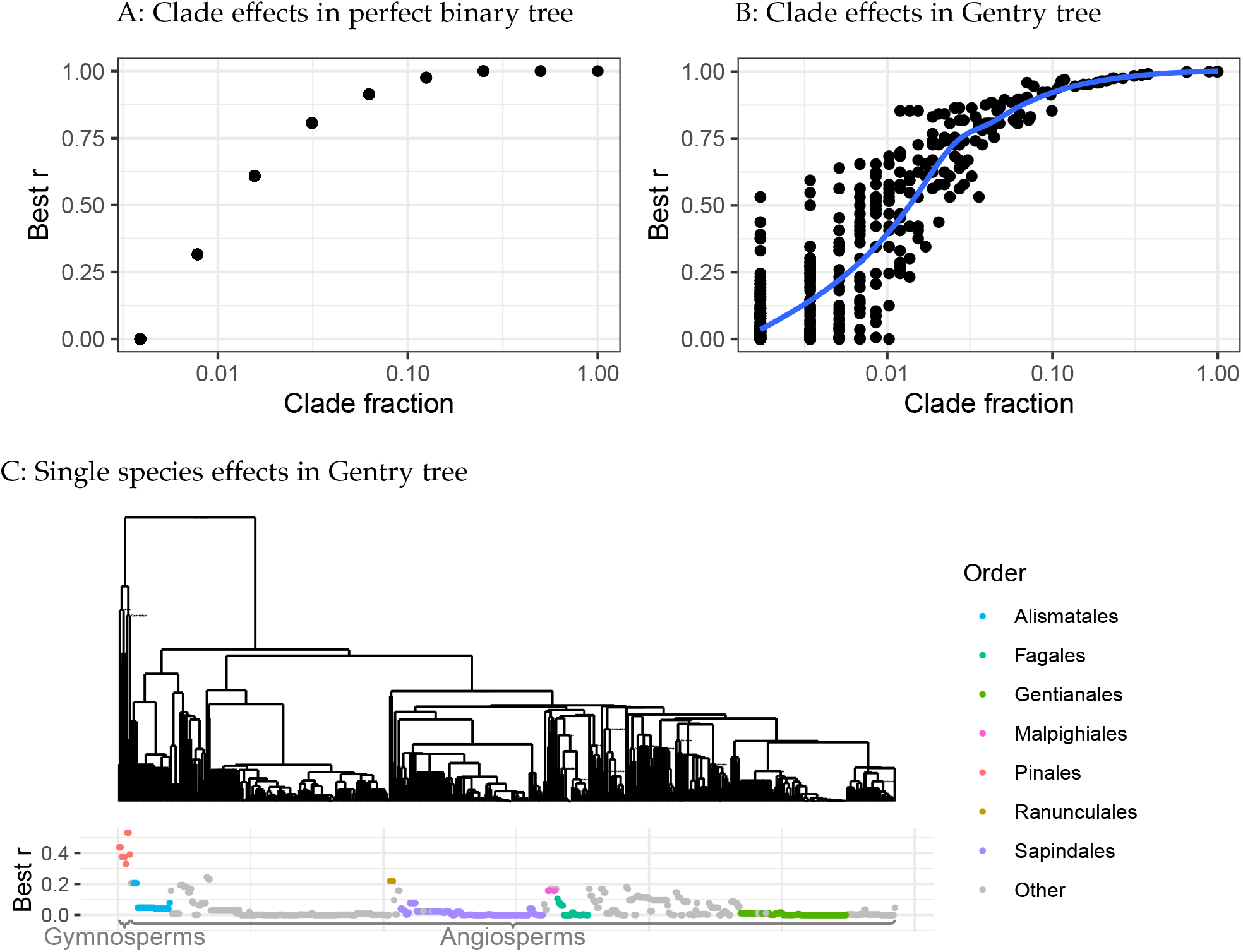
The best value of *r* for detecting an effect that is constrained to one clade is dependent on the size of that clade. (A) The best value of *r* for all possible “clade effects” in the perfect binary tree. Each point corresponds to a clade, the position on the *x*-axis shows the fraction of the total number of leaves in the tree the clade represents, and the position on the *y*-axis shows the best value of *r* for detecting a difference in that clade. (B) Analogous to A for the phylogenetic tree corresponding to the Gentry dataset. (C) Best value of *r* for all possible “single species effects” in the Gentry tree. Top of panel shows the Gentry dataset phylogeny. Each point on the bottom panel shows the best value of *r* for detecting a difference between sites when the difference is confined to a single species. Points are aligned to the phylogenetic tree, and *y*-axis position gives the best value of *r* for detecting the difference.

In other words, the two communities have the same abundance for all species aside from those in one clade, which are uniformly over-represented in site 1. We then compute the best value of *r* for detecting a difference between samples when it is of the form given above. The results are shown in Figure 4A, where each point in the figure represents one clade. The position of a point on the horizontal axis represents the size of that clade as a fraction of the total number of species in the tree, and the position of the point on the vertical axis shows the value of *r* that gives us maximal power to see a difference in that clade. We see that when the difference is in individual species (a clade fraction of 1/256 ≈ .004), values of *r* close to 0 are best, but when the effect is in large clades (clade fractions above .1), values of *r* close to 1 are best.

This effect holds true for real trees as well. In Figure 4B, we see the same computation, but in this case the tree is taken from the Gentry dataset (described in the Section 4.2 below), and ***µ***_*x*_ and ***µ***_*y*_ differ in one clade in that tree as described above. As before, each point in the figure represents one clade, the position of a point on the horizontal axis represents the size of that clade as a fraction of the total number of species in the tree, and the position of the point on the vertical axis shows the value of *r* that gives us maximal power to see a difference in that clade.

As with the perfect binary tree, when ***µ***_*x*_ and ***µ***_*y*_ differ in a large clade, larger values of *r* perform the best, while when they differ in a small clade, smaller values of *r* perform best. However, although the general shape of the relationship is similar in the real tree and the perfect binary tree, there is much more variability in the best value of *r* for a given clade size, particularly for the small clades. This is due to the fact that there is more heterogeneity in the relationships among the species in the real tree than in the perfect binary tree.

To get more insight into what is driving variability in the best value of *r*, we can focus on the singleton “clades” (i.e. individual species), corresponding to the points at the far left of Figure 4B. The modal best value of *r* for detecting differences between sites that are confined to an individual species (“best *r* for singleton clades”) is 0 (101 of 585 singletons, or about 17% had 0 for the best value of *r*). The mean best value of *r* is similarly small, at about .04. However, there are several species for which the best value of *r* is quite high, over .25, which was never the case for the perfect binary tree. If we look into this more carefully, we see that the species for which the best value of *r* is highest are those species that are quite different from the rest A: Clade effects in perfect binary tree of the tree. We see this in Figure 4C, where we have computed the best value of *r* for detecting a difference confined to each of the individual species in the Gentry dataset. The phylogenetic tree corresponding to the Gentry dataset is plotted at the top of Figure 4C, and the best value of *r* for detecting a difference when ***µ***_*x*_ and ***µ***_*y*_ differ in only one species is plotted below the phylogeny. We see that the species corresponding to very large best values of *r* are those that are quite different from the rest of the tree. In particular, the all the species with the highest best value of *r* are those that are on the left side of the most basal split in the dataset. These correspond to the Gymnosperms in the dataset, while all the other species are Angiosperms.

## 4 Examples

So far, we have shown theoretically how the PQ distance can be decomposed into parts related to basal vs. terminal phylogenetic structure, that it generally places a large amount of weight on the basal structure, and that we can get more power to detect effects at different phylogenetic scales by modifying the weights. We show next how the standard PQ distance doesn’t have the power to see real phylogenetically-structured effects and how altering the skewed weight distribution can help. We see this both in simulated and real data.

### 4.1 Simulated example

We will first show how we can use the MPQ_*r*_ distances in simulated data. We will set up the simulation to illustrate how different members of this family emphasize different sorts of patterns. To that end, we will imagine that we are sampling communities from sites that vary in two dimensions, e.g., we are looking at sites in a mountain range, and the sites vary in both their altitude (high or low altitude) and average moisture content of the soil (wet or dry sites). In the simulation, there will be two true differences in species abundance: the first will be that a large clade is consistently under-represented at high sites and over-represented at low sites. The second true difference will be that there are two sister clades, one of which is over-represented in the wet sites the other of which is over-represented in the dry sites. Furthermore, there will be some negative phylogenetic autocorrelation (if a species is over-represented at a site, its closest relatives will tend to be under-represented at that site), and altitude and moisture will be roughly uncorrelated with each other.

The probabilistic model corresponding to this description is the following:

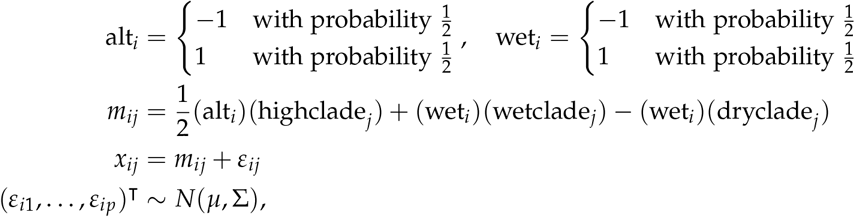

where alt_*i*_ = 1 means site *i* is at high altitude, alt_*i*_ = −1 means that site *i* is at low altitude, wet_*i*_ = 1 means site *i* is wet, and wet_*i*_ = −1 means site *i* is dry. highclade_*j*_ = 1 if *j* is a species in the clade associated with altitude and 0 otherwise, wetclade_*j*_ = 1 if *j* is a species in the random set of species positively associated with the wet sites gradient and zero otherwise, and dryclade_*j*_ = 1 if *j* is a species in a random set of species negatively associated with wet sites and zero otherwise.

To create highclade, wetclade, and dryclade, we created a random tree with 300 leaves using the `rtree` function in `ape` [34]. For highclade, we took a node that has 132 descendants and let highclade_*j*_ = 1 for each of those 132 descendants. For wetclade and dryclade, we took a node that had 47 descendants. One of the children of that node had 27 descendants, and wetclade_*j*_ = 1 for each of those 27 descendants. The other child of that node had 20 descendants, and dryclade_*j*_ = 1 for each of those 20 descendants.

Σ is chosen so that there is some negative phylogenetic autocorrelation. All the code for the simulation is in the `simulations` vignette in the `mpqDist` package so that users can experiment with different settings for these parameters.

In Figure 5, we see the differences between the two types of sites as quantified by different MPQ_*r*_ distances. In each panel, we compute MPQ_*r*_ distances between each site and a reference site (one of the low/wet sites). As we might expect, low and wet sites are always closer to the reference site (which is low and wet) than high or dry sites are. However, the amount of separation between the two classes we see is strongly dependent on *r*. For the altitude effect, which is associated with the large clade, we see that the difference between low and high sites is the clearest for *r* = 1, intermediate for *r* = .5, and hardly visible at all for *r* = 0. In contrast, the moisture effect, which is associated with the two smaller clades, is the clearest for *r* = .5, intermediate for *r* = 1, and again hardly visible at all for *r* = 0.

**Figure 5:**
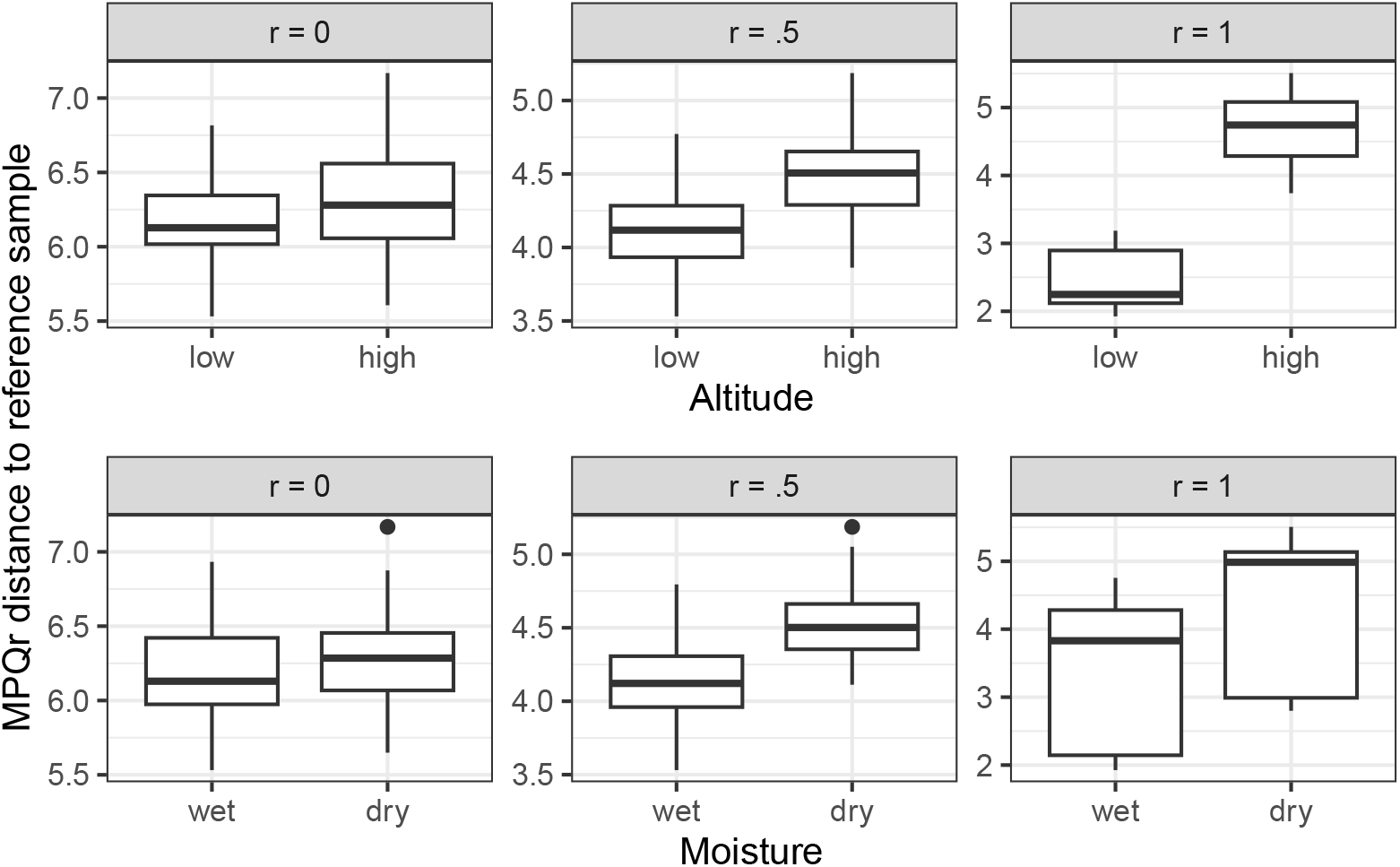
Large values of *r* are more effective for the altitude effect and moderate values are more effective for the moisture effect. Each boxplot shows the distribution of MPQ_*r*_ distances between each site and a reference site. Each facet is labeled with the value of *r* used for the MPQ_*r*_ distance. Sites are separated out into low- and high-altitude sites in the top row and into wet and dry sites in the bottom row.

The simulation illustrates two things. First of all, when there is a phylogenetic structure to the data, we tend to get more power (in the statistical sense) by using a phylogenetically-informed distance (MPQ_*r*_ with *r >* 0). Secondly, the different vaules of *r* preferentially emphasize effects on different phylogenetic scales. In this case, *r* = 1 shows the effect associated with a large clade the most clearly, while *r* = .5 shows the effect associated with an intermediate-sized clade the most clearly. In this particular simulation, if we had used only the Euclidean distance (MPQ_*r*_ with *r* = 0) to look for the altitude/moisture effects, we would have concluded that there was very little difference between the sites. If we had used only the PQ distance (MPQ_*r*_ with *r* = 1), we would have concluded that there was an altitude effect but very little moisture effect. By using the full range of values of *r* in the MPQ_*r*_ family, we can see both of true effects, and we can also correctly infer that the moisture effect is associated with a finer phylogenetic scale than the altitude effect.

### 4.2 Latitudinal species turnover

To illustrate how these distances can be used to further our understanding of species turnover, we look at the Gentry forest transect dataset [39]. The dataset gives the abundances and identities of all plants with stem diameters at breast height of more than 2.5cm in ten 2 × 50 meter transects at each of 197 sites. The sites span six continents, although they are not evenly distributed, with the western hemisphere and sites near the equator being more common. The raw data are available on the Missouri Botanical Garden website [1], and as has been done in previous studies [10], the localities used in this study were taken from the `anadat-r` website [2].

A phylogenetic tree was created for the species using S.Phylomaker [42]. As S.Phylomaker uses taxonomic information to build the phylogenetic tree, a fair amount of cleaning was necessary to extract the information necessary to build the tree. Interested readers can see the code in the `gentry.R` file in the `mpqDist` package, but the main operations were to remove patterns in the name such as “cf,” “aff,” “var,” and so on, so that tentative species or variants are collapsed into a single species. In addition, there were several genera whose corresponding family has been changed (e.g. Azara used to be in the family Flacourtiaceae and is now classified as Salicaceae). When these were identified by S.Phylomaker, we updated the family name. The process resulted in less than 2% of the counts in the original dataset corresponding to species that S.Phylomaker could not assign to a location in the phylogeny. These were dropped from the analysis.

We looked at the species turnover from high to low latitudes using several MPQ_*r*_ distances. We took as a reference the northern temperate sites (between 23.5 and 66.5 degrees north). For each site, we calculated the average MPQ_*r*_ distance between that site and the set of northern temperate sites for *r* = 0 (Euclidean distance), *r* = 1 (PQ distance), and an intermediate value of *r*. The sitewise abundances were all transformed according to a centered log-ratio transform [3] before computing these distances (see Appendix A.5, particularly the last paragraph, for why transformation is reasonable). To help interpret these results, we compared them to three other methods of quantifying species turnover. We computed Bray-Curtis distances (without transforming the species abundances) to allow us to compare to a standard non-phylogenetic measure of dissimilarity. We also agglomerated species to either the order level or the division level and then calculated the Euclidean distance between the centered log-ratio transformed agglomerated counts. The idea here was to give more intuition about the taxonomic levels that correspond to different values of *r* in the MPQ_*r*_ distances.

In Figure 6, we see plots of the average dissimilarity between each site and the northern temperate sites as measured by each of the six distances we are comparing. Each point represents a site, and the error bars represent the mean plus or minus two standard errors for each fifteen-degree-of-latitude bin. Starting with the PQ distance (*r* = 1), the plot primarily shows us that the equatorial sites (around latitude = 0) are farther from the temperate sites than the 40 degree samples. There is a suggestion that the southern temperate sites (circa 40 degrees south) are farther from the northern temperate sites than the equatorial sites are, but the effect is not very clear, and there are quite a few equatorial sites that are farther from the northern temperate sites than any of the southern temperate sites are according to this measure. Finally, the very most northern sites, those around 60 degrees north, are also farther on average from the northern temperate sites, but there are very few of them.

**Figure 6:**
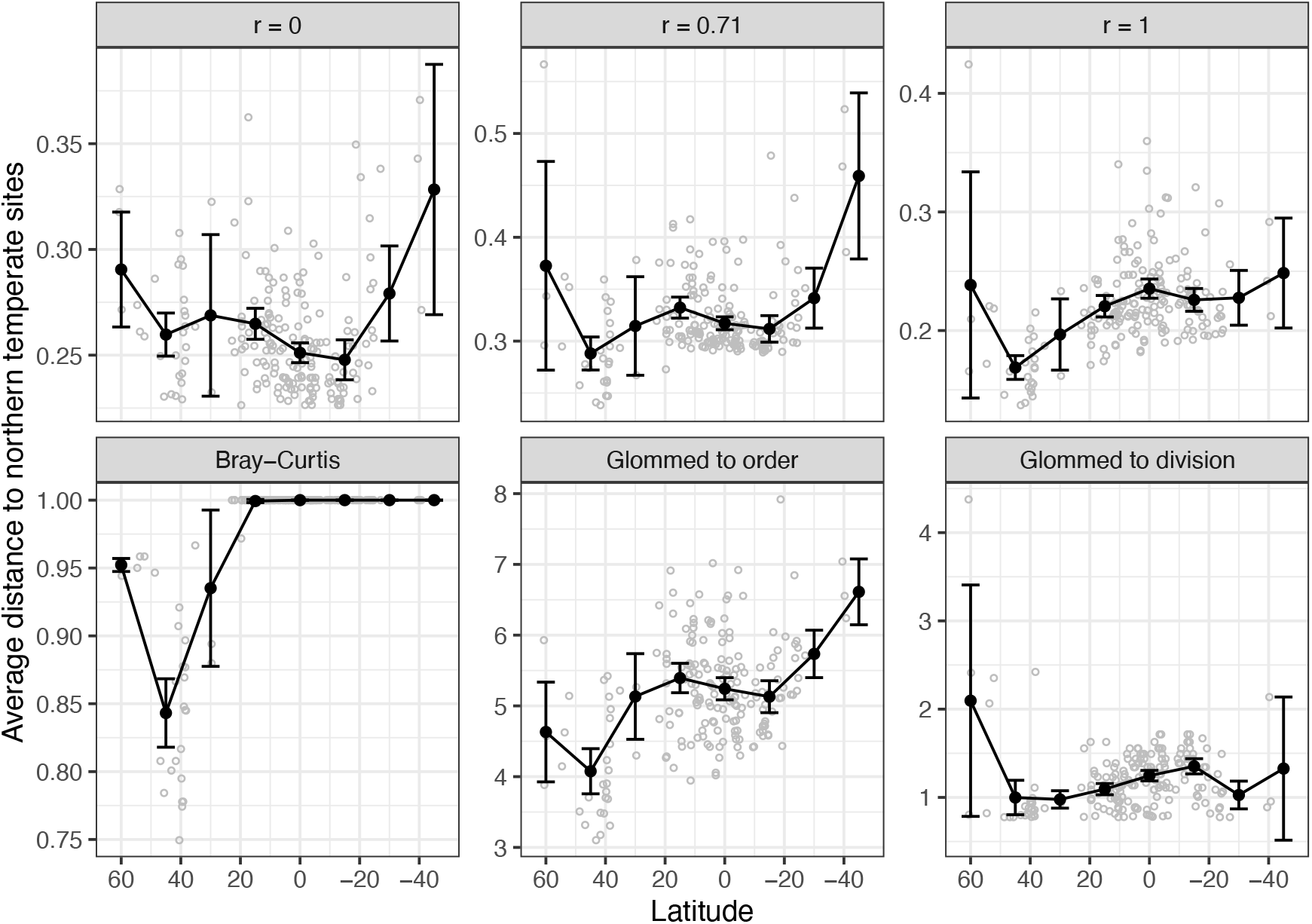
Average distance of each site to the northern temperate sites, as measured by six different distances. Each gray circle corresponds to one site. Position of the point on the *x*-axis represents its latitude. Position on the *y*-axis represents the average distance between that site and the set of northern temperate sites (sites between 23.5 and 66.5 degrees north). Black points and error bars represent the average distance in a 15-degree of latitude bin plus or minus two standard errors. Different facets represent different distances: The top row is MPQ_*r*_ distances with *r* = 0 (Euclidean distance), *r* = .71 (intermediate between Euclidean and PQ), and *r* = 1 (PQ distance). Bottom row, left to right, is Bray-Curtis dissimilarity, Euclidean distance on species agglomerated to the order level, and Euclidean distance an species agglomerated to the division level.

In contrast, the MPQ_*r*_ distance with *r* = .71 shows more dynamic range. As with *r* = 1, the equatorial sites are more dissimilar from the northern temperate sites than the northern temperate sites are from each other. However, unlike with *r* = 1, we see another uptick at the farthest southern sites, with the 40 degrees south sites being more dissimilar from the northern temperate sites than the equatorial sites are.

Finally, the MPQ_*r*_ distance with *r* = 0 (the Euclidean distance) has some strange behavior that illustrates why it is not commonly used for ecological data analysis. According to the Euclidean distance, the very most northern and very most southern sites are the most dissimilar to the northern temperate sites. However, it also suggests that the 10 degrees south sites are more similar to the northern temperate sites than the 20 degrees north sites are, or even than those sites are to themselves. This apparent similarity is not due to overlap in species composition, rather, it is due to the fact that the 10 degrees south sites have uniformly fewer species in them. There is therefore a larger amount of overlap between the species that are absent at the 10 degrees south sites and the northern temperate sites than there is between the species that are absent at the 20 degrees north sites and the northern temperate sites. This is of course not what we are trying to measure, and is a large part of the reason why Bray-Curtis is used for comparing species composition instead of the Euclidean distance.

Therefore, as another point of comparison, we look at the performance of the Bray-Curtis dissimilarity on the same task. As we see on the bottom row of Figure 6, the issue with Bray-Curtis is that it saturates. There is no overlap between the species at the northern temperate sites and the species present below 20 degrees north, and so the Bray-Curtis dissimilarity is 1 for all save one of the sites below 20 degrees north and the northern temperate sites. It does, however, do a better job of putting the sites above 20 degrees north in the proper order of dissimilarity than the Euclidean distance.

Finally, we look at distances computed on the agglomerated species counts. When we use Euclidean distance on counts at the order level, the qualitative effect is much like the MPQ_*r*_ distance with the intermediate value of *r*, where the equatorial sites are farther from the northern temperate sites and the southern temperate sites are farther than that. Similarly, the Euclidean distance on counts at the division level (which, as a reminder, is very coarse for this dataset, with only two divisions represented), we see that the qualitative effect is more similar to the MPQ_*r*_ distance with *r* = 1. There is a slight increase in distance going from the northern temperate sites to the equatorial sites, not a clear difference between the southern temperate sites and the equatorial sites, and a very large difference between the northern temperate sites and the sites around 60 degrees north.

## 5 Data and code availability

We have created an R package that allows users to easily compute PQ and MPQ_*r*_ distances. It integrates with `phyloseq` [33], which is the most streamlined way of storing the combined species abundance and phylogenetic tree data structure that is required for this kind of analysis. It also enables users to easily create animated/interactive plots where users can scroll through the complete range of values of *r* in the MPQ_*r*_ distances. The package is available on github https://github.com/jfukuyama/mpqDist and can be installed using `devtools::install_github(jfukuyama/mpqDist)`. The package includes a vignette descrbing the overall functions and vignettes that reproduce the simulated and real data examples in Sections 4.1 and 4.2. The R script at https://github.com/jfukuyama/mpqDist/blob/main/data-raw/gentry. R gives the code used to clean the Gentry dataset.

## 6 Discussion

We showed that the PQ distance places an overwhelming amount of weight on derived variables related to the basal structure of the phylogeny. To do so, we derived an interpretation of the PQ distance as a weighted Euclidean distance on derived variables related to the species abundances and the phylogeny, showed what those derived variables were, and derived the weights in some special cases. We also showed that a second class of distances, the MPQ_*r*_ distances, have the same interpretation but place less weight on the basal phylogenetic structure. Our power results show that the standard PQ distance has the most power to detect differences between communities that are related to the basal phylogenetic structure of the tree (for example, if the difference between communities is primarily at the phylum or division level). For differences between communities that are related to more terminal phylogenetic structure (for example, differences between sites that are confined to smaller clades or taxonomic groups), analysts can gain statistical power by using one of the MPQ_*r*_ distances or another phylogenetically-informed distance that places less emphasis on basal structure. Our analysis of the Gentry transect data suggested that the PQ distance emphasizes very broad-scale differences among communities: the PQ distances among sites looked the most like distances computed on division-level informtion about species composition of the sites.

These results sharpen the existing literature on phylogenetically-informed measures of beta diversity. There have been empirical demonstrations that phylogenetically-informed beta diversity measures can be classified into terminal and basal measures [45, 20]. Swenson [45] classifies Rao’s D [43] and pairwise phylogenetic dissimilarity [46] as basal measures, whereas UniFrac and PhyloSor [8] are terminal measures. There has also been empirical work suggesting that comparing the results of terminal vs. basal measures can give insight into the scale of phylogenetic scale at which community structure varies with environmental factors [26]. Our work shows the mathematical basis for the terminal-to-basal gradient in one class of phylogenetically-informed distances, and our power analysis results give a theoretical justification for the practice of comparing terminal and basal measures.

Although our results give more insight into the PQ and MPQ_*r*_ distances, there is room for improvement. We have argued that the weight accorded to the basal phylogenetic features by the PQ distance is too high and that the MPQ_*r*_ distances modulate the amount of weight accorded to them in a resonable way. However, that is not the only way to do the modulation, and it is possible that other strategies are better. In addition, we have argued that different values of *r* in the MPQ_*r*_ distances are analogous to considering only information at different taxonomic ranks, but the correspondence is not transparent and probably varies depending on the particulars of the set of species.

Since we do not have a good interpretation of different values of *r* past the observation that larger values of *r* give a more “basal” phylogenetic measure, it is hard to give recommendations about what specific values of *r* to choose. One reasonable way to proceed is to perform the analysis using a variety of different values of *r*. This is analogous to recommendations in the literature to perform analyses with different beta diversity measures in order to better understand the dataset at hand [5], for instance, measures based on presence/absence vs. relative abundance vs. abundance (e.g. Jaccard, Sorensen vs. chi squared vs. Euclidean), measures that include or exclude joint absences (e.g. Jaccard vs. simple matching), and different transformations of abundances to prior to computing standard measures (all reviewed in [5]). It is also related to the recommendations to compare the results of terminal and basal phylogenetic beta diversity measures described above. The `mpqDist` package has this sort of functionality, allowing users to create interactive plots based on MPQ_*r*_ distances that let the user scroll through the range of values of *r*. This sort of interactive plot is similar in spirit to diversity profiles that emphasize rare vs. common species [28].

One potential issue with this strategy is that introducing a family of distances gives more “researcher degrees of freedom” [23], and can lead to invalid *p*-values if the value of *r* is chosen based on looking at the data and the data-driven choice of *r* is not taken into account when computing *p*-values. There are several potential ways to deal with this, all of which are standard for situations in which there are a large number of different ways to analyze a particular dataset. First of all, the method and results can be purely exploratory, with no attempt at statistical inference/*p*-value computation. If the analyst wants a *p*-value based on an MPQ_*r*_ distance, they could do something similar to a traditional power analysis, where they simulate data that has the signal and noise structure that they expect, use the simulated data to choose the value of *r* that will have the most power, and use that value of *r* to analyze their experimentally-generated data. If the analyst does not have a good sense of what he expects the real signal to look like, he could also split the data into two pieces, using the first piece to decide on a good value of *r* and the second piece to compute *p*-values.

We hope that the results presented here give ecologists a better understanding of the measures available for characterizing dissimilarities between communities and allow them to gain more insight into the biological processes controlling community composition.

## A Appendix

### A.1 Equivalent and closely related measures

In the following sections, we list some of the related measures and describe their connection to the PQ distance. The PQ distance is based on Rao’s axiomatization of diversity. His framework is both extremely influential and extremely flexible, which means that the PQ distance is either closely related to or the same as several other measures that have been described in the literature.

#### A.1.1 Chave, Chust, and Thébaud’s *H*

Chave, Chust, and Thébaud [12] describe a phylogenetically-informed dissimilarity measure between two communities that they call *H*. They suppose that the divergence time between taxon *i* and taxon *j* is denoted by *t*_*ij*_. They assume that the *t*_*ij*_’s are scaled so that the maximal divergence time is 1, and then set *P*_*ij*_ = 1 − *t*_*ij*_ if *i* ≠ *j* and *P*_*ii*_ = 1. Notice that this means that their *P*_*ij*_ values are the same as the elements of **T** for an ultrametric tree (that is, a tree for which the distance from each leaf to the root is the same) for which the distance from root to tip is one. They then define 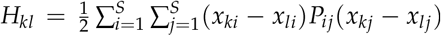. If we let **P** be a matrix whose *ij*th element is *P*_*ij*_ and **x**_*k*_ = (*x*_*k*1_ *x*_*k*2_ … *x*_*kS*_)^⊺^, then the matrix form of this equation is 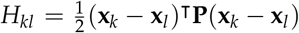. Since we have already seen that **P** is a special case of a similarity or covariance matrix **T**, as defined in Section 2.2, we see that 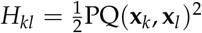.

#### A.1.2 Distance between communities in double principal coordinates analysis

Double principal coordinates analysis is an ordination method that gives a low-dimensional representation of a set of communities and the species that make up those communities. It is also based on Rao’s framework for diversity. As the exact mathematical description of DPCoA requires a lot of notation, we give a high-level overview of the procedure here. We will use the same notation as the original paper describing DPCoA [37] so that interested readers can connect the description here to the description in the paper.

Assume we have the abundances of *n* species in *r* communities, and we have a *n × n* matrix **Δ**_*n*_ that contains dissimilarities among the species. This **Δ**_*n*_ is not the same as the one used in the main section of the paper. For DPCoA generally, it can be anything and we are later going to specialize it to be square roots of patristic distances on the tree. To perform DPCoA, we perform the following steps:

1. Using principal coordinates analysis, create a matrix **X ∈** *R*^*n*×*n*−1^ such that the distances between the rows of **X** match the distances in **Δ**_*n*_.
2. Create a matrix **Y ∈** ℝ^*r*×*n*−1^. Each row of **Y** corresponds to one community such that each community is placed at the barycenter of its species points.
3. Finally, perform PCA on **Y** to get a low-dimensional representation of the sites. Since the species and sites are represented in the same *p −* 1-dimensional space in (2), the species points in **X** can be projected onto the principal axes to help aid interpretation.

The authors note that if we let 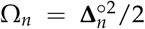 (where ∘2 indicates element-wise square of the matrix) and if we let **p**_*i*_ and **p**_*j*_ be the vectors containing the relative abundances of the species in sites *i* and *j*, then

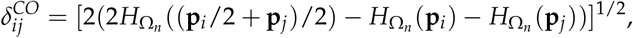

where 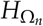 in the DPCoA paper is the same as the 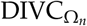 function defined in Section 2.1. They also note that 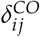 is the distance between sites *i* and *j* in the matrix **Y** described in point (2) above. Therefore,

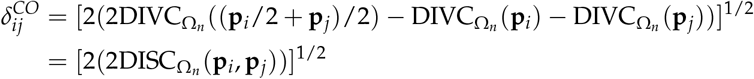

Therefore, if we use the square roots of the patristic distances between the species in DPCoA (that is, if the matrix **Δ**_*n*_ described in the second paragraph of this section contains the square roots of the patristic distances among the species), then 2Ω_*n*_ will be a matrix with the patristic distances between the species. Since 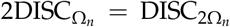, this means that 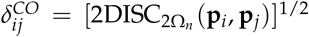. Since 2Ω_*n*_ contains patristic distances, this means that 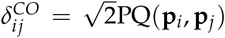. In particular, this means that the distances between the communities in **Y** from point (2) above is equal to 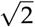 times PQ(**p**_*i*_, **p**_*j*_).

There is an extra factor of 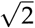 because in Pavoine et al. define DISC_**Δ**_(**p**_1_, **p**_2_) = 2DIVC_**Δ**_((**p**_1_ + **p**_2_)/2) − DIVC_**Δ**_(**p**_1_) − DIVC_**Δ**_(**p**_2_) (Equation (2) in [37]), while we follow Rao’s original definition of 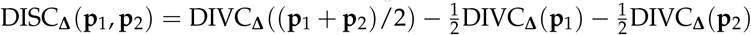 (Equation 2.1.3 in [43]).

The switch from the patristic distances used in the PQ distance to their square roots might seem puzzling at first, but it is actually the correct choice in the context of DPCoA. This is because DPCoA requires an embedding of the species in Euclidean space in such a way that the distances between the embedded points match the input distances. The square roots of the patristic distances are always Euclidean [13], while the raw patristic distances generally are not (with the exception of ultrametric trees).

#### A.1.3 Hardy and Senterre’s *P*_*ST*_

Hardy and Senterre [24] have a measure of phylogenetic beta diversity they call *P*_*ST*_. The focus of this paper is on distances, but any measure of beta diversity can be converted to a measure of dissimilarity between simply by defining the dissimilarity between two communities to be the beta diversity of the set containing those two communities. Using the same **Δ** matrix as in Section 2.1 (that is, **Δ** contains the patristic distances between the species), supposing that **p**_1_, … , **p**_*m*_ are vectors representing species proportions at each of *m* locations, and letting 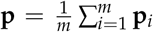, they define

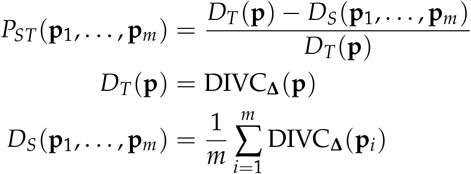

From this, we see that *P*_*ST*_(**p**_1_, **p**_2_)*D*_*T*_(**p**) = *D*_*T*_(**p**) − *D*_*s*_(**p**_1_, **p**_2_) = DISC(**p**_1_, **p**_2_) = PQ(**p**_1_, **p**_2_)^2^.

### A.2 Maximal value of PQ

#### Theorem A1.

*Suppose* **Δ** *is a p × p matrix corresponding to the patristic distances among the p leaves of a tree. Let i*_1_, *i*_2_ *be a pair of leaves such that*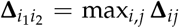. *Then* 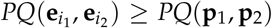 *for any* **p**_1_, **p**_2_ *such that* 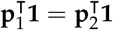 *and* **p**_1_, **p**_2_ ⪰ 0.

*Proof*. Recall that 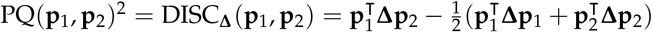.

We can take the three terms independently. First, consider 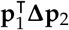. The elements of **Δp**_2_ must all be between 0 and 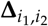 because each element of **Δp**_2_ is a convex combination of the elements of **Δ**. Similarly, 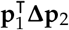 must also be between 0 and 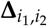 because it is a convex combination of the elements of **Δp**_2_, which we just established were all between 0 and 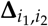 . Since 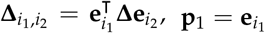 and 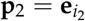 must be the maximizing values of 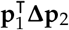.

Next, notice that 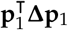 is a weighted sum of the elements of **Δ**, all of which are at least zero. It therefore has a minimum value of 0, and 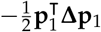 has a maximum value of 0. Since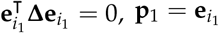, maximizes 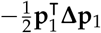.

The same holds for 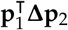 and 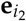 , and so we can see that PQ(**p**_1_, **p**_2_) takes its maximal value when 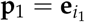 and 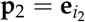 . □

### A.3 Eigendecomposition of the perfect binary tree similarity/covariance matrix

In this section, we derive the eigenvectors and eigenvalues of the matrix **T** described in Section 2.2 corresponding to the perfect binary tree with 2^*K*^ leaves. The proof relies on the fact that this matrix can be written as a sum of block diagonal matrices. In the main text, the eigenvectors are the coefficients of linear derived variables and help us interpret the PQ distance. However, since **T** is a covariance matrix, the eigenvectors we derive here can also be interpreted as the principal components of **T**.

#### Theorem A2.

*Suppose* **T**^*pbt*^ *is the similarity/covariance matrix described in Section 2.2 corresponding to a perfect binary tree with* 2^*K*^ *leaves and edges with length 𝓁*_*k*_ *at level k, k* = 0, … , *K* − 1. *Assuming that the species are arranged as they are in Figure 1A, the first eigenvector of* **T**^*pbt*^ *is*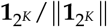, *and the subsequent eigenvectors are of the form* 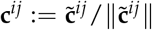, *with*

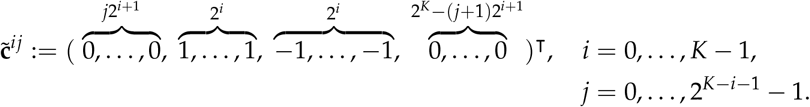

*The eigenvalue corresponding to the eigenvector* 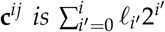 , *and the eigenvalue corresponding to* 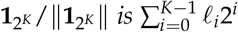.

*Proof*. Let **B**^*i*^ be a 2^*K*^ × 2^*K*^ with 2^*i ×*^ 2^*i*^ blocks of 1’s on the diagonal. The matrix **T**^pbt^ corresponding to the tree can be written as

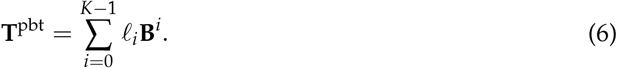

Notice that every **c**^*ij*^ is an eigenvector of every **B**^*i′*^ . We have the following relation:

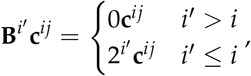

and so **c**^*ij*^ are eigenvectors with eigenvalue 0 of **B**^*i′*^ if *i*′ *> i* (the block in the **B**^*i′*^ is bigger than the block in **c**^*ij*^) and are eigenvectors with eigenvalue equal to the size of the block in **B**^*i′*^ otherwise.

Substituting this result into equation (6), we see that the **c**^*ij*^ are the eigenvectors of **T**^pbt^, and

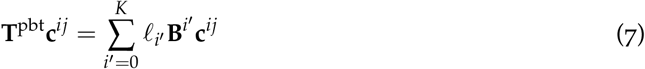

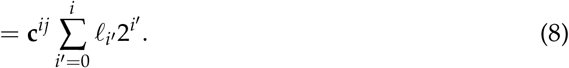

Therefore, every **c**^*ij*^ is an eigenvector of **T**^pbt^, and the eigenvalue corresponding to **c**^*ij*^ is 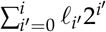 Similarly, **B**^*i*^**1** = 2^*i*^**1**, which implies that

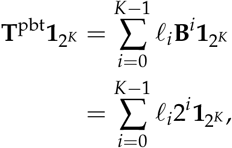

and so 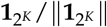 is an eigenvector of **T**^pbt^ with eigenvalue 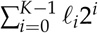. □

**Note A1**. *From line (8), we see that* **c**^*ij*^ *with larger values of i (i*.*e*., *non-zero on larger clades) will always have larger eigenvalues than those with smaller values of i (non-zero on smaller clades)*.

**Note A2**. *The eigenvectors have a simple interpretation:* **c**^*ij*^ *corresponds to a difference in average values between sister clades of size* 2^*i*^. *The normalization ensures that features corresponding to different clade sizes are on the same scale — without such normalization, the features corresponding to larger clades would in expectation be larger simply because they correspond to sums over larger collections of variables. Finally, the eigenvalue corresponding to an eigenvector of the form* **larger eigenvalues than those with smaller valuecs**^*ij*^ *will be* 2^*i*+1^ − 1.

**Note A3**. *Supposing that 𝓁*_*i*_ = 1 *for all i (all the branch lengths are equal to 1), the sum of the eigenvalues corresponding to eigenvectors of the form* **c**^*K*−*i*,*j*^ *is* 2^*K*^ − 2^*K*−*i*−1^. *Therefore, the sum of all the eigenvalues is*

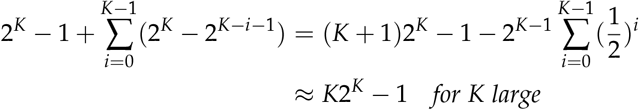

*Since the largest eigenvalue is* 2^*K*^ − 1, *this implies that the first eigenvalue contributes approximately* 1/*K of the overall trace*.

**Note A4**. *To aid intuition, we can write down what all these matrices are for a small perfect binary tree. Let* **T**^*pbt*^ *be the perfect binary tree with four leaves and with branch lengths all equal to* 1/2, *so that the distance from the root to each leaf is 1. Then*

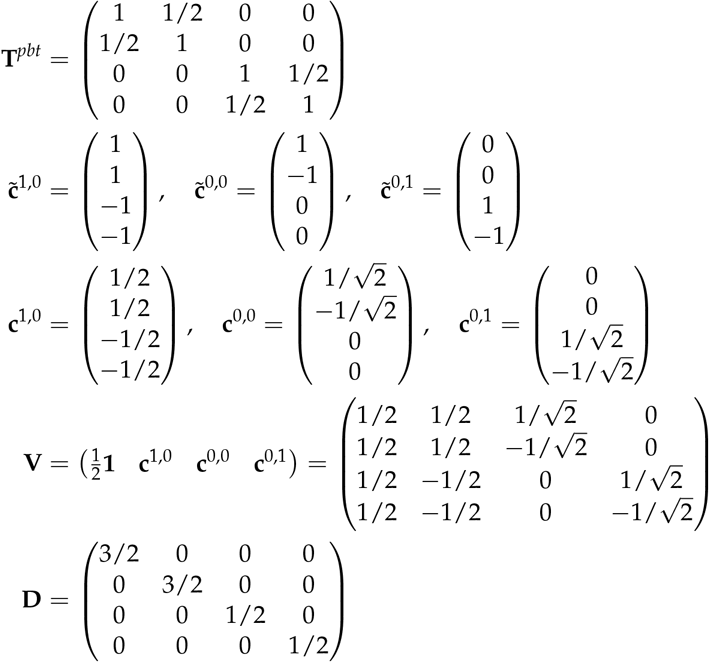

### A.4 Eigendecomposition of the comb tree similarity/covariance matrix

In this section, we derive approximations to the eigenvectors and eigenvalues of the matrix **T** described in Section 2.2 corresponding to a comb tree. Before we give the main result, we describe the comb tree we will analyze and the matrix **T** that corresponds to it.

The comb tree we will be considering is a binary tree. An example with 5 leaves is illustrated in Figure 7. In the general case with *p* leaves, we will let *l*_0_, … , *l*_*p*−1_ denote the *p* leaves and *n*_0_, … , *n*_*p*−2_ denote the *p* − 1 internal nodes. *n*_0_ is the root. There are 2(*p* − 1) edges in the graph, two for each internal node. There are edges between *n*_*i*_ and *l*_*i*_, *i* = 0, … , *p* − 2, edges between *n*_*i*_ and *n*_*i*+1_, *i* = 0, … , *p* − 3, and an edge between *n*_*p*−2_ and *l*_*p*−1_. All edges aside from the one connecting *n*_*p*−2_ and *l*_*p*−1_ have length 1/*p*. The edge connecting *n*_*p*−2_ and *l*_*p*−1_ has length 2/*p*. Note that the tree is scaled such that the distance between the root and leaf *l*_*p*_ is equal to 1, and that the matrix describing the amount of shared ancestral branch length between nodes can be written as

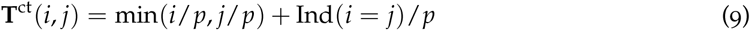

**Figure 7:**
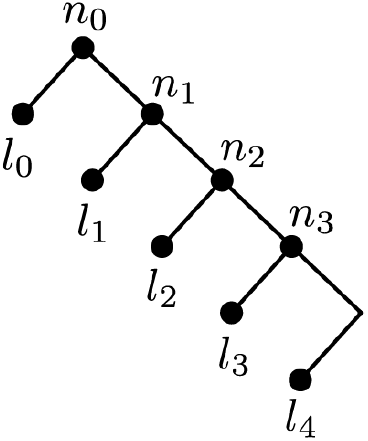
A diagram showing a comb tree with five leaves and four internal nodes.

We will only be able to get approximate results for the comb tree. The lemma below gives the exact result that the approximations will approach for bigger and bigger comb trees. *K* plays the role of **T** with *K*(*x, y*) being analogous to one of the elements of **T**.

#### Lemma A1.

*Let K*(*x, y*) = *min*(*x, y*), *for* (*x, y*) ∈ [0, 1] × [0, 1]. *The eigenfunctions of K are f*_*a*_(*y*) = sin(*yaπ*/2) *for a an odd integer, and the eigenvalue corresponding to f*_*a*_(*y*) *is* 4/(*π*^2^*a*^2^).

*Proof*.

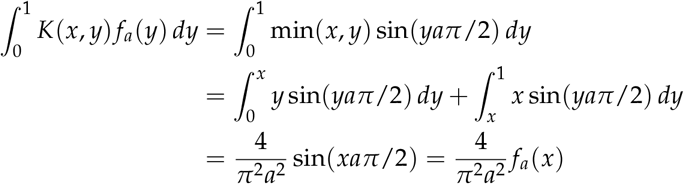

□

#### Theorem A3.

*Let* **T**^*ct*,*p*^ = **T**^*ct*^/*p, a scaled version of the proximity matrix defined in equation (9) Let f*_*p*,*a*_(*i*) = sin(*aπi*/2*p*), *for a an odd integer. Then*

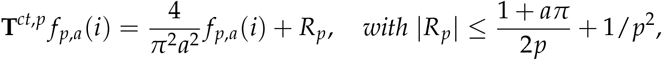

*meaning that f*_*p*,*a*_ *is an approximate eigenvector of* **T**^*ct*,*p*^. *Additionally*,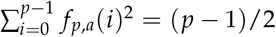 *except when a*/(2*p*) ∈ ℤ, *and so the normalized approximate eigenvectors will be* 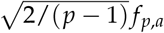.

*Proof*. We can write

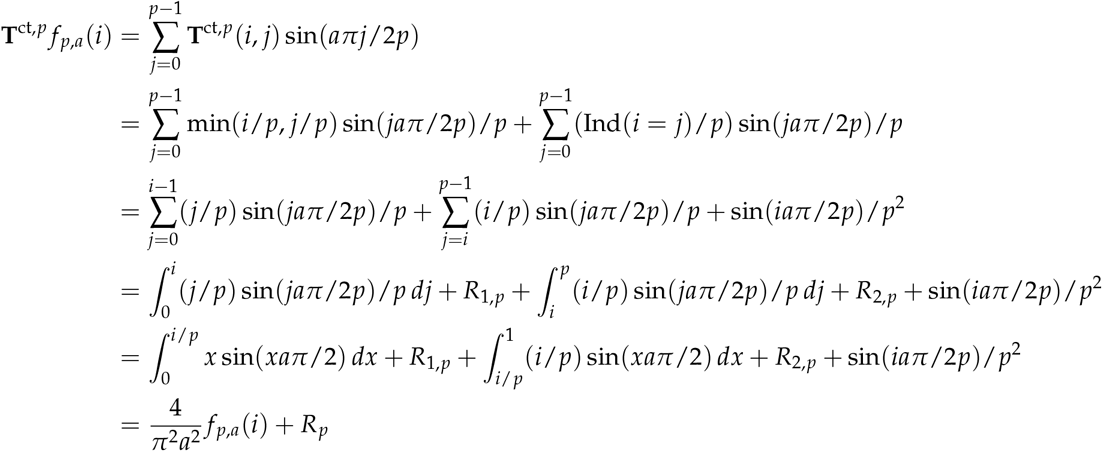

where *R*_*p*_ = *R*_1,*p*_ + *R*_2,*p*_ + sin(*iaπ*/2*p*)/*p*^2^, and *R*_1,*p*_ and *R*_2,*p*_ are the error bounds for numerical integration using the left-hand rule and the last line follows by the lemma and because *f*_*p*,*a*_(*i*) = *f*_*a*_(*i*/*p*). *R*_1,*p*_ and *R*_2,*p*_ satisfy

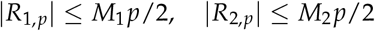

for

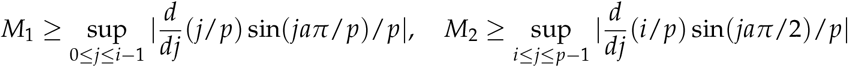

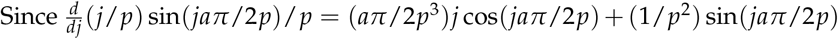 and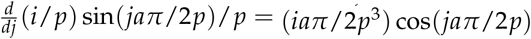 , we can take *M*_1_ = (1 + *aπ*/2)/*p*^2^ and *M*_2_ = *aπ*/2*p*^2^.

Then if we set *R*_*p*_ = *R*_1,*p*_ + *R*_2,*p*_ + sin(*iaπ*/2*p*)/*p*^2^, we see that as required.

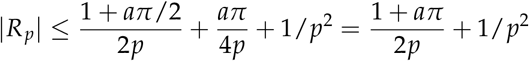

□

**Note A5**. *The sum of the eigenvalues of K can be written as* 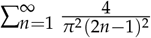, *which converges and is equal to* 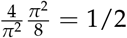. *The fraction of the trace contributed by the first eigenvalue is therefore* 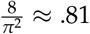.

### A.5 Interpretation of MPQ_*r*_ distances as distances between smoothed matrix estimates

We showed in the main text that the MPQ_*r*_ distances allow us to modulate the weights in the PQ distance. However, there are a large number of different ways that one could in principle modulate the weights, and we would like to have a reason for choosing this one. One motivation for using the scheme defined by the MPQ_*r*_ distances is that the MPQ_*r*_ family has an interpretation in terms of a Bayesian model for the species abundances that incorporates the phylogenetic information as a prior.

Suppose we let **x**_*i*_ be a *p*-dimensional vector of observations corresponding to the *i*th sample. Our model for **x**_*i*_ is as follows

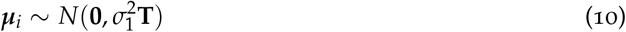

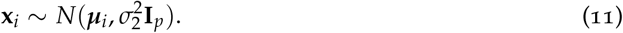

The idea is that we believe the true means, given in ***µ***_*i*_, to be informed by the phylogenetic tree. The data we observe, **x**_*i*_, is a corruption of that signal with spherically-symmetric normal noise. In this model, we would like to know about the true means, ***µ***_*i*_, and so the natural quantity to compute is the posterior distribution of ***µ***_*i*_ given **x**_*i*_. This is

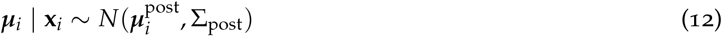

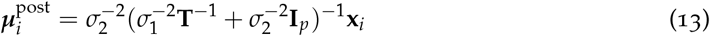

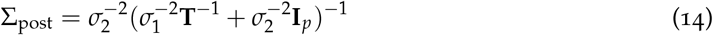

Recall that the Mahalanobis distance can be used to measure the difference between two points when they are assumed to come from multivariate normal distributions with different means but the same variance. In particular, if **x**_1_ and **x**_2_ come from multivariate normal distributions with common covariance matrix Σ, the Mahalanobis distance between them is [(**x**_1_ − **x**_2_)^⊺^Σ(**x**_1_ − **x**_2_)]^1/2^ [31]. The posteriors have common covariance, and so we can measure distances between the posterior means, taking into account the posterior variance, by using the Mahalanobis distance with Σ_post_ as the covariance matrix:

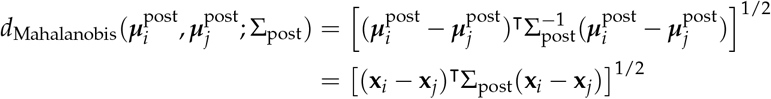

Then, since 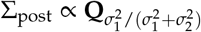, we see that the Mahalanobis distance [31] on the posterior means is the same, up to a scaling factor, as Equation (1).

Note that the model in (11) and the posterior distribution in (12) are analogous to Gaussian process regression. In Gaussian process regression, the assumption is that you have noisy observations of a function that is smooth in some space, and the smoothness assumption is enforced by putting a Gaussian process prior on the noise-free observations. In model (11), we are assuming that we have noisy observations of a true mean vector that is smooth on the tree, and we enforce the smoothness assumption by using **T** as the covariance for the prior on the true means. In particular, the model in (11) and the posterior in (12) is analogous to the Gaussian process regression model described in Rasmussen and Williams [48] (our posterior is a slightly rearranged version of the posterior in equations (2.22)-(2.24) in Rasmussen and Williams). Our model is analogous to their model in Chapter 2, with our **x**_*i*_ playing the role of their **y**, our ***µ***_*i*_ playing the role of their **f**, and our **T** playing the role of their kernel function *K*. This model was also used before for analyzing microbiome data [19], although without the Gaussian process description.

Some implications of this interpretation are as follows:

– The distances given by the MPQ_*r*_ distances can be interpreted as the distances between rows of the “smoothed” or “denoised” data matrix.
– The standard Euclidean distance (equivalent to MPQ_0_) is the limit as the assumed observation error goes to 0.
– The PQ distance (equivalent to MPQ_1_) is the limit as the assumed observation error goes to ∞.
– The normal-model explanation of the distances suggests that these methods will work best when the data can be reasonably expected to come from a signal plus normal noise model.

The last point suggests that raw counts or proportions, which we do not expect to come from something close to a normal distribution, will not be accounted for well by this model. This problem is consistent with our experience using these distances: we get better results from both the PQ distance and the MPQ_*r*_ distances when we work with counts that have been put through a variance-stabilizing transformation instead of raw counts or proportions, for instance, using log-ratio or centered log-ratio transformations of the original data as was done in Section 4.2.

### A.6 Generalized *χ*^2^ distribution parameterizations

#### Definition A1

(Standard parameterization of generalized *χ*^2^ distribution). *The standard way of parameterizing the generalized χ*^2^ *distribution is to say that* 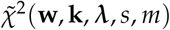 *has the distribution of* ∑_*i*_ *w*_*i*_*χ*^′2^(*k*_*i*_, *λ*_*i*_) + *sz* + *m, where χ*^′2^(*k*_*i*_, *λ*_*i*_) *are independent non-central chi-squared random variables with degrees of freedom k*_*i*_ *and non-centrality parameters λ*_*i*_ *and z is an independent standard normal random variable*. □

The distributions we will use here will be more naturally written as quadratic forms of multivariate normal random variables, and so it is worthwhile to write down the correspondence between the quadratic form representation and the standard parameterization.

#### Lemma A2.

*Suppose* **z ∼** *N*(*µ*, Σ). *Let* Σ^1/2^ *be a symmetric square root of* Σ, Σ^1/2^**Q**Σ^1/2^ = **VDV**^⊺^ *be the eigendecomposition of* Σ^1/2^**Q**Σ^1/2^, *and* ***ν*** = (**V**^⊺^Σ^−1/2^*µ*)^°2^ (**x**^°2^ *indicates the element-wise or Hadamard square of the vector* **x***). Then*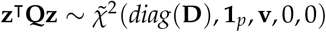.

*Proof*. Let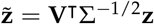. Then 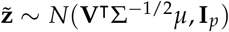, and we can write

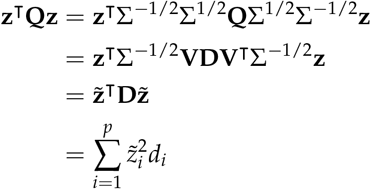

where 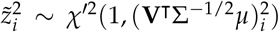, where *χ*^′2^(*a, b*) denotes a non-central chi-square distribution with degrees of freedom *a* and non-centrality parameter *b*. Therefore, 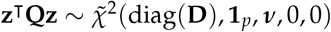 in the notation of Definition 1. □

### A.7 Justification of scaling

#### Theorem A4.

*Suppose* **x, y ∼** *N*(**0**_*p*_, **I**_*p*_) *and are independent of each other. Then E*[*MPQ*_*r*_(**x, y**)^2^] = 2. *In particular, the expected squared distance does not depend on r*.

*Proof*. For any value of *r*, we can write **Q**_*r*_ = **VD**_*r*_**V**^⊺^, where **V ∈** ℝ ^*p*×*p*^ is orthonormal and **D**_*r*_ ∈ *ℝ*^*p*×*p*^ is diagonal with positive elements. Since **V** is orthonormal, 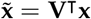 and 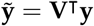 are distributed *N*(**0**_*p*_, **I**_*p*_), still independently of each other. With that knowledge, we can write

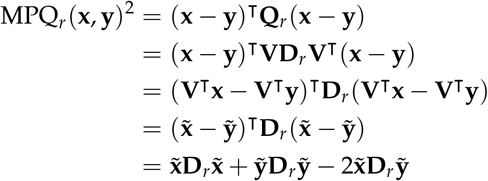

Then notice that

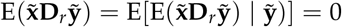

and that 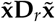 and 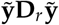 are each distributed 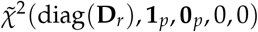. The means are therefore 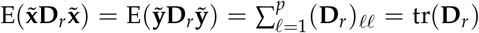. When we constructed **D**_*r*_, we normalized it so that its trace was 1 , and so putting the three pieces together tells us that E _*r*_ [MPQ (**x, y**)^2^] = 2 for any value of *r*. □

### A.8 Power analysis

We would like to understand how a statistical test based on the MPQ_*r*_ distance would perform for different sorts of signal and different sorts of noise. We will define what the test would be and write down an expression for its power. We will also derive an approximation to the test threshold that will give us more insight into the regimes in which tests based on MPQ_*r*_ with small vs. large values of *r* are likely to perform well.

#### Definition A2

(MPQ_*r*_ test). *Let* **x, y** ∈ ℝ^*p*^ *have distribution* **x** ∼ 𝒩 (*µ*_*x*_, Σ) *and* **y** ∼ **𝒩** (*µ*_*y*_, Σ). *We assume* Σ *is known and that we have specified a fixed value of r. The null hypothesis is H*_0_ : *µ*_*x*_ = *µ*_*y*_.

*The MPQ*_*r*_ *test rejects when MPQ*_*r*_(**x, y**)^2^ *falls above the* 1 − *α quantile of a generalized chi-squared distribution with parameters* **w** = *Evals*(2Σ^1/2^**Q**_*r*_Σ^1/2^), **k** = **1**_*p*_, ***λ*** = **0**_*p*_, *s* = 0, *m* = 0.

#### Lemma A3

(MPQ_*r*_ test has the correct size). *Under the null hypothesis H*_0_ : *µ*_*x*_ = *µ*_*y*_, *the probability that the MPQ*_*r*_ *test will reject H*_0_ *is equal to α*.

*Proof*. Under the null hypothesis, **x** − **y** ∼ *N*(**0**_*p*_, 2Σ). Therefore, by Lemma 2, (**x** − **y**)^⊺^**Q**_*r*_(**x** − **y**) ∼ 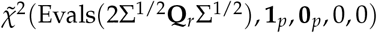, and so a test that rejects according to the 1− *α* quantile of this distribution will reject with probability *α*. □

#### Lemma A4

(Power of MPQ_*r*_ test). *Supposing that µ*_*x*_ ≠ *µ*_*y*_, *the power of the MPQ*_*r*_ *test is given by*

*P*(*S* ≥ *q*_1−*α*_), *where S is a generalized chi-squared random variable with parameters* **w** = *Evals*(2Σ^1/2^**Q**_*r*_Σ^1/2^), **k** = **1**_*p*_, ***λ*** = **V**^⊺^Σ^−1/2^(*µ*_*x*_ *− µ*_*y*_), *s* = 0, *m* = 0, *q*_1−*α*_ *the upper quantile of the distribution described in Definition 2*.

*Proof*. In this case, **x** − **y** ∼ *N*(*µ*_*x*_ − *µ*_*y*_, 2Σ), and so again by Lemma 2, 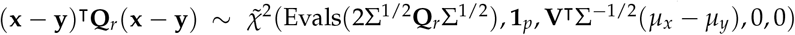. □

## Notes

### Competing Interest Statement

The authors have declared no competing interest.

